# Sequencing of a dairy isolate unlocks *Kluyveromyces marxianus* as a host for lactose valorization

**DOI:** 10.1101/2025.02.03.636257

**Authors:** Mackenzie Thornbury, Adrien Knoops, Iain Summerby-Murray, James Dhaliwal, Sydney Johnson, Joseph Christian Utomo, Jaya Joshi, Lauren Narcross, Gabriel Remondetto, Michel Pouliot, Malcolm Whiteway, Vincent J.J. Martin

**Author notes:** Swiss Federal Institute of Technology Lausanne (EPFL), Rte Cantonale, 1015 Lausanne, Switzerland. Department of Chemistry, University of Alberta, 11227 Saskatchewan Drive, Edmonton, Alberta, Canada T6G 2G2 (Joseph Christian Utomo). Department of Wood Science, University of British Columbia, Forest Sciences Centre, 2424 Main Mall #2900, Vancouver, BC V6T 1Z4.

## Abstract

The use of genetically modified non-conventional yeast provides significant potential for the bioeconomy by diversifying the tools available for the development of sustainable and novel products. In this study, we sequenced and annotated the genome of *Kluyveromyces marxianus* Y-1190 to establish it as a platform for lactose valorization. The strain was chosen for rapid growth on lactose-rich dairy permeate, high transformation efficiency, and ease of culturing in bioreactors. Genomic sequencing revealed that *K. marxianus* Y-1190 possesses single nucleotide polymorphisms associated with efficient lactose metabolism. The strain is diploid with notable genomic heterogeneity, which appears to be critical for its robust growth and acid tolerance. To further exploit this platform strain, we developed protocols for gene and chromosome manipulation using CRISPR editing, constructed and validated a series of promoters compatible with MoClo vectors, and designed synthetically inducible promoters for *K. marxianus*. These tools enable precise control over gene expression, allowing for the tailored optimization of metabolic pathways and production processes. The synthetic promoters provide flexibility for dynamic expression tuning, while the CRISPR-based editing protocols facilitate targeted genetic modifications with high efficiency. Together, these advancements significantly enhance the genetic toolbox for *K. marxianus*, positioning it as a versatile platform for industrial biotechnology. These tools open new opportunities for the sustainable production of bio-based chemicals, fuels, and high-value products, leveraging lactose-rich feedstocks to contribute to a circular economy.

## INTRODUCTION

The search for efficient and cost-effective hosts for industrial applications is an ongoing process in biotechnology. *Kluyveromyces marxianus*, a non-conventional yeast species, has gained considerable attention in recent years due to specific properties that make it a suitable candidate for a variety of industrial applications^1^. *K. marxianus* is known for its high growth rate^2^, together with broad pH tolerance^3^, heat tolerance^4^, and ability to metabolize a wide range of sugars^5,6^. These characteristics make it an ideal candidate for use in industrial processes such as bioethanol production^7–9^, the production of value-added products such as enzymes^10–12^, and bioactive compounds^13,14^. *K. marxianus* strains have been isolated from a variety of sources including dairy products, fermented foods, and soil^6^. Several strains of *K. marxianus* have been studied for their potential use in industrial applications. Among the commonly studied strains for industrial applications are NBRC1777 and DMKU3-1042, isolated from soil in Thailand and Japan, respectively, as well as CBS 6556, which was isolated from the fermented drink pozol^15–17^. Although these strains can grow on lactose as a carbon source, they are not from the dairy lineage of *K. marxianus*^18^ and may not be suited for applications in the dairy industry.

Despite being produced in great quantities as a by-product of dairy processing, lactose is the weak link of the modern dairy industry. All other outputs of dairy processing – cream, skim milk, cheese and protein concentrates – sell into markets that are consistently profitable. In contrast, the lactose byproduct of dairy processing, referred to as permeate, is a low value commodity that in North America is sold into highly volatile markets. In many cases the available price for lactose does not justify the cost of concentrating and drying this by-product for sale as animal feed, and these lactose streams must be disposed of at an environmental and economic cost. Milk and whey permeates are often used as cheap options for food sweeteners, however, with the rate of lactose intolerance and the popularity of lactose-free products, that market for dairy permeate may be shrinking^19^. Developing markets for the lactose fraction of non-fat solids is a longstanding issue that the dairy industry has been grappling with for years.

Advances in genetic engineering and the development of molecular tools have paved the way for the manipulation of this yeast species to improve its industrial potential. To date, two promoter libraries^20,21^, a genetic toolkit^22^, and standard protocols for CRISPR have been developed for *K. marxianus*^23–25^. While these tools have been instrumental in broadening the field to work with *K. marxianus,* few are standardized and the majority work with non-dairy isolates that can respire but not ferment lactose, rather than dairy isolates that can do both^26^.

In the current study, we aimed to domesticate a strain of *K. marxianus* capable of transforming the low-value lactose found in dairy permeate into higher-value specialty chemicals. We screened 19 strains of *K. marxianus* available from public collections and identified six candidates with robust growth on both minimal lactose medium and lactose-rich dairy permeate. We further screened the 19 strains on milk permeate with fumaric acid added to pH 2.2, since for bioprocessing, limiting the addition of a base during fermentation can streamline the process and cut down costs^27^. Many molecules of commercial interest are acidic, such as the nine organic acids that have been designated by the United States Department of Energy as chemical building blocks that can be produced from biomass^28,29^. Having a strain that can be leveraged to metabolize lactose and be organic acid-tolerant would be an asset to the circular economy. We tested the plasmid transformation efficiency of the characterized strains from these screens, identifying three strains that grew well in all conditions and had high transformation efficiencies.

We have generated the first publicly available annotated genome of the lactose-permeate-using *K. marxianus* strain Y-1190. We have further modularized and characterized 23 promoters sourced from a promoter library^20^ and the *K. marxianus* toolkit^22^ for activity on lactose, glucose, and milk permeate in both strain Y-1190 and a soil strain CBS 6556. Additionally, we took three of the strongest promoters and made them tetracycline inducible. We further developed an improved CRISPR-Cas9 system for genomic deletions and integrations using a combination of double-guide array and an NHEJ inhibitor and finally, we optimized the fermentation strategy for Y-1190 in a bioreactor to ensure its industrially relevant potential as a production strain.

## RESULTS

### *K. marxianus* strain selection

To identify a strain of *K. marxianus* that not only grows well on lactose, but also grows well on dairy permeate, a substrate that differs in composition from commonly used laboratory media, we compared the growth rate of 19 *K. marxianus* strains grown in supplemented dairy permeate with those grown in YNB lactose (YNBL) medium (Fig 1A and B). We identified six strains that grew at a rate above 0.4 1/h in both media: Y-2415, Y-1190, Y-12992, Y-12990, Y-1163, and L388 (Fig 1A).

**Figure 1.**
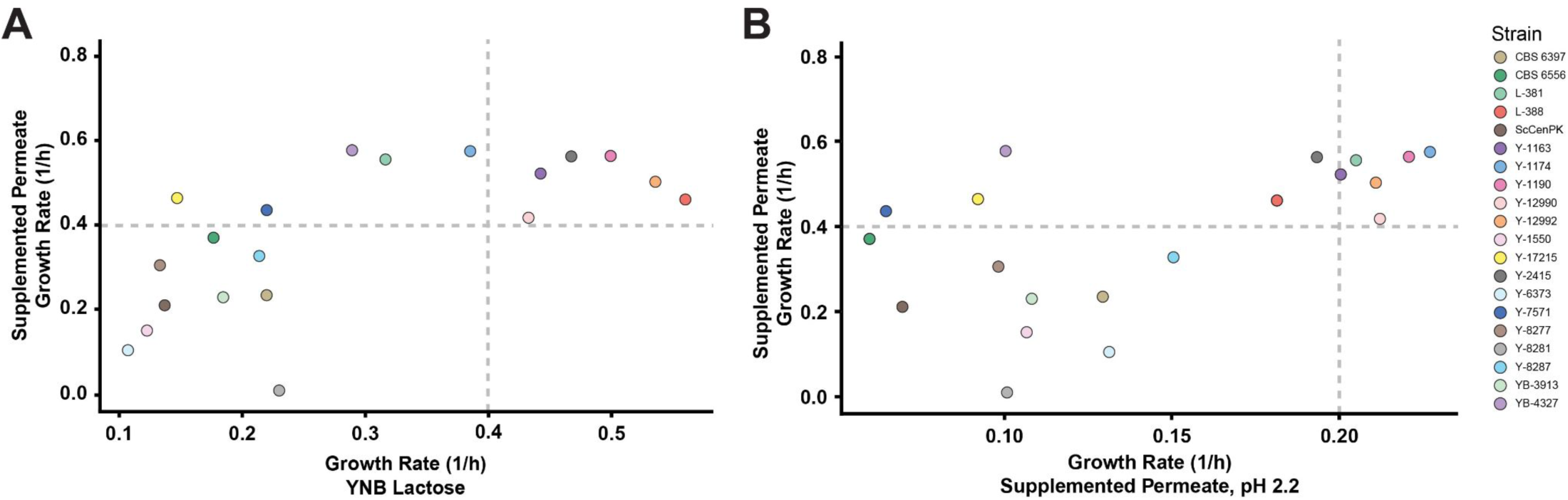
Growth rate comparison of aerobic growth of 19 *K. marxianus* strains on **A**. YNBL and Supplemented Permeate and **B**. Supplemented Permeate and Supplemented Permeate at pH 2.2.

Additionally, organic acid tolerance is an important phenotypic trait from a bioprocessing standpoint. We compared the growth of our strains in supplemented permeate without adjusting the pH (pH 6) and at an acidic pH of 2.2 by adding fumaric acid to its solubility limit of 5 g/L (Fig 1B), and identified six strains (Y-1190, Y-12990, Y-12992, Y-1163, L381, and Y-1174) that grew above 0.4 1/h for the supplemented media and 0.2 1/h for the supplemented permeate at pH 2.2 (Fig 1B). The raw growth rates for all strains in YNBL, YNBG and supplemented permeate can be found in the supplemental data (Fig S1).

### Transformation efficiency

To identify isolates that are most genetically tractable, we tested DNA transformation efficiency of the six strains selected for robust growth on dairy permeate and low pH. The strains were transformed with a modified Gietz procedure using an empty vector plasmid containing a spacer and nourseothricin resistance^30^. From this data, three strains, Y-1190, Y-12990, and Y-12992, had comparable transformation efficiencies of 1.14×10^4^, 1.44×10^4^, and 2.38×10^4^ cfu/ug DNA respectively (Fig S2), and surpassed our growth rate thresholds in all three media conditions (Fig. 1). We selected strain Y-1190 based on these data in combination with previously published work that highlighted its genetic tractability and high transformation efficiency^23^.

### Genomic sequence analysis

We created a publicly available annotated genome for *K. marxianus* strain Y-1190 to facilitate the use of this yeast in strain engineering projects. Strain Y-1190 was sequenced by PacBio technology, which generated two assembly files: one primary assembly and one secondary assembly containing haplotigs, establishing the diploid nature of the strain. The primary assembly resulted in 10 contigs and the secondary assemblies resulted in 66 haplotigs. To confirm the near-chromosome level of our assemblies, each contig was queried using NCBI BLAST while restricted to sequences from *K. marxianus* (taxid 4911). Primary contigs were found to be close to chromosome length and total assembly size matched closely to the *K. marxianus* reference strain DMKU3-1042 – 1.1×10^7^ for Y-1190 compared to 1.09×10^7^ in DMKU3-1042 (Table S1).

Before gene prediction, low complexity and highly repetitive regions were masked using RepeatMasker (see Methods). Repetitive sequences made up 245885/10951256 bp (2.25 %) while low complexity sequences made up 39868/10951256 bp (0.36 %). Using AUGUSTUS, gene prediction resulted in the identification of 4853 putative genes for Y-1190 with an average gene length of 1549 nt. Gene prediction for the haplotig data identified 4108 genes. The primary assembly was classified as 98.08 % complete single-copy, 0.09 % as complete duplicated, 0.28 % as fragmented, and 1.54 % as missing. In the haplotig assembly 76.28 % were classified as complete single-copy, 1.59 % as complete duplicate, 0.89 % as fragmented, and 21.24 % as missing. When predicting genes by BUSCO, missing genes could mean that there was no significant hit in the database for that gene. This is highly possible since there are only two *K. marxianus* genomes in the BUSCO database, both non-dairy strains^31^.

#### Analysis of lactose permease variation

Previous work has identified three groups of *K. marxianus* haplotypes, A, B and C, which differ on their ability to metabolize lactose^26^. While all three haplotypes can metabolize lactose, only haplotype B can both ferment and respire lactose. This is thought to be due to polymorphisms in the lactose permease Lac12-1, which increased lactose transport 10 fold in dairy isolate CBS 397 compared to non-dairy isolate CBS 6556^32^. With full access to the Y-1190 genome, we conducted a multisequence alignment of *LAC12-1* genes from Y-1190, dairy isolate CBS 397, and non-dairy isolates CBS 6556, NBRC 1777, and DMKU3-1042 (Fig 2). Our analysis revealed that the *LAC12-1* genes of Y-1190 and CBS 397 exhibit 19 identified polymorphisms, aligning with those observed in the Varela *et al.* study, and identifies the *LAC12-1* gene in Y-1190 as encoding the more active variant (Fig 2).

**Figure 2.**
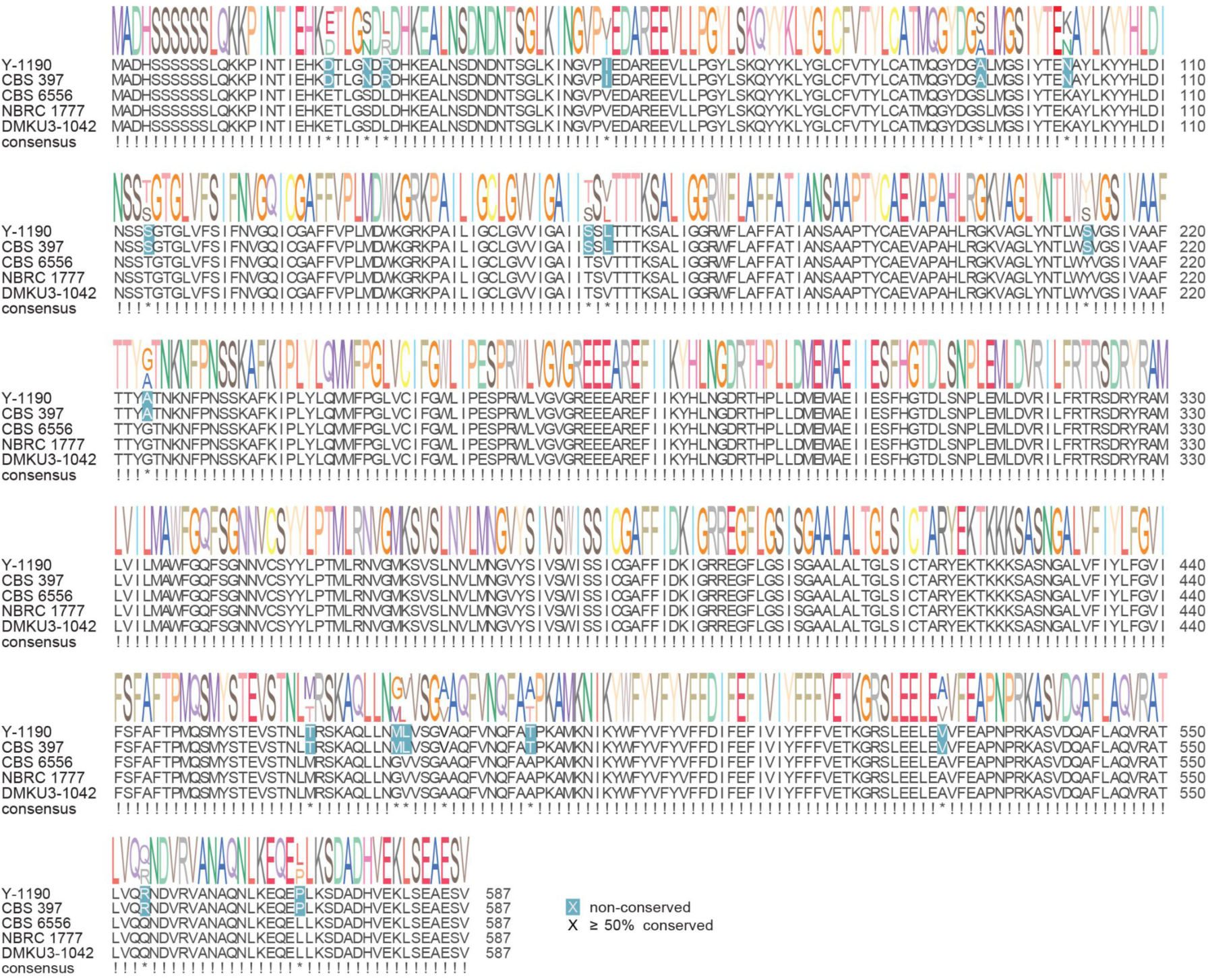
Comparison of *LAC12* genes in *K. marxianus* strains Y-1190, CBS 397, CBS 6556, NBRC 1777, and DMKU3-1042.

### Promoter set characterization

To allow efficient engineering of recombinant gene expression in *K. marxianus* Y-1190, we characterized the activity of a collection of promoters of varying strength in cultures grown on lactose-containing media. Previously, three groups have characterized *K. marxianus* promoter libraries^20–22^. These works focused on glucose and xylose metabolism, and many of the promoters tested were not standardized. As part of our characterization of lactose metabolizing strain *K. marxianus* Y-1190, we modified 21 promoters from Lang *et al.*^20^ to adhere to the modular cloning standard described for *Saccharomyces cerevisiae*^33^ and tested them alongside two other promoters from Rajukumar *et al.*^22^ for a total of 23 promoters (Table S2).

Our standardized promoter library was cloned upstream of sfGFP and was transformed into two strains of *K. marxianus*: the non-dairy strain CBS 6556 and the dairy strain Y-1190. The library was tested in YNBL, YNB glucose (YNBG) and supplemented permeate (Fig 3, Fig S4). To ensure the design of the constructs would work both in lactose and permeate, we compared the promoter strengths of those two conditions (Fig 3A). Generally, the relationship is linear, with promoters tending to drive overall higher expression in supplemented permeate than YNBL, apart from *pTDH* which performs 4.3-fold better in YNBL than supplemented permeate. Similarly, the relationship between promoter expression in YNBL and YNBG is linear with the only exception being *pLAC4* which is only expressed in YNBL, as expected (Fig 3B). After characterizing the promoters in strain Y-1190 across different media, we compared the promoter expression profiles between Y-1190 and the soil-derived strain CBS 6556 in both YNBL (Fig 3C) and YNBG (Fig 3D). There were several promoters that fell outside a strictly linear relationship. As an example, *pINU1* (light orange) consistently had higher expression in Y-1190 while *pRPE1* (light brown) consistently had higher expression in CBS 6556 regardless of the carbon source (Fig 3CD). Both p*PDC1* (medium blue) and *pNC1* (medium green) were consistently high for both strains in all conditions, although with differing magnitudes of expression (Fig 3CD). All expression data for strains CBS 6556 and Y-1190 in YNBL, YNBG, and supplemented permeate at 6 hrs and 24 hrs can be found in the supplemental data (Fig S4).

**Figure 3.**
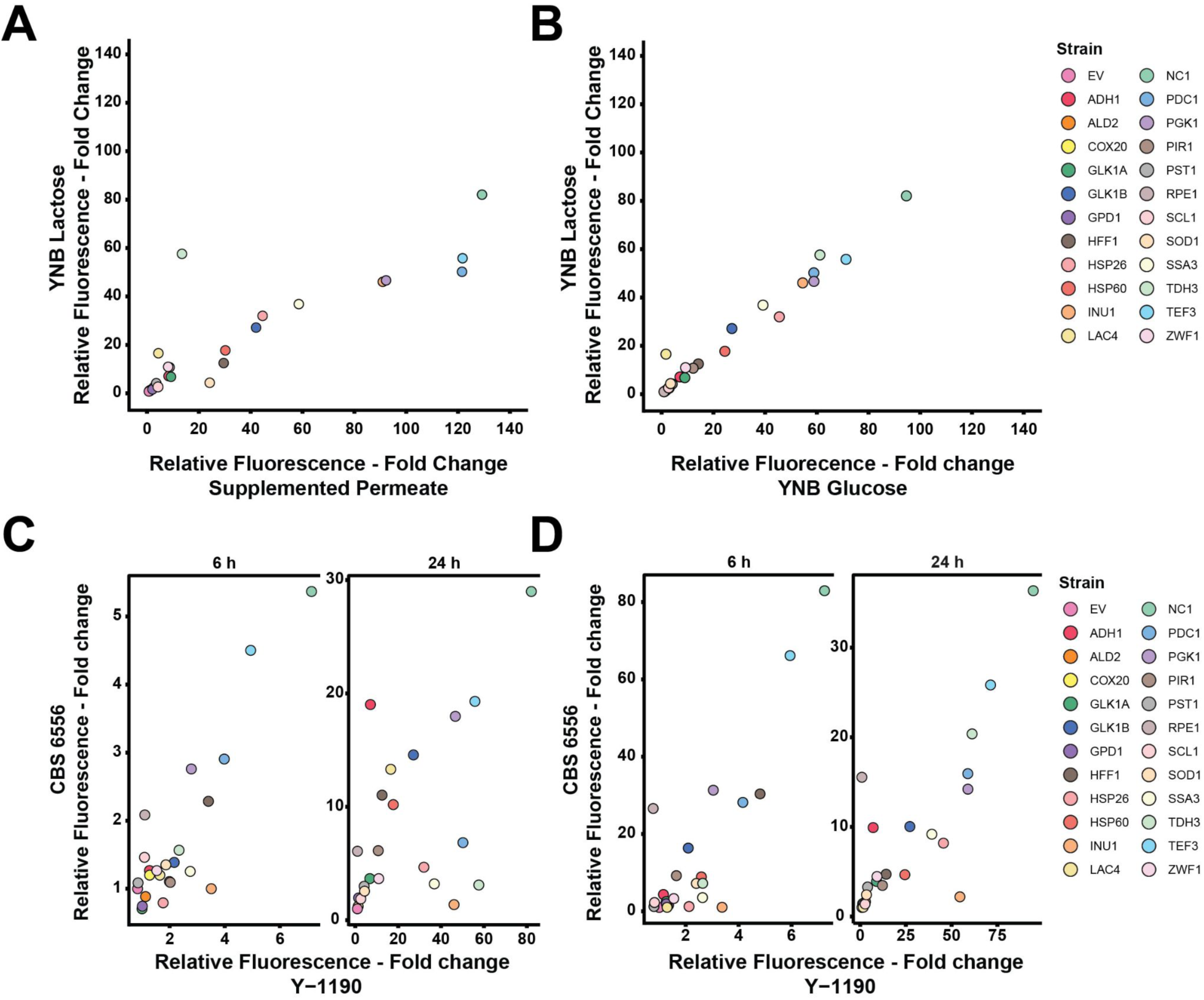
Characterization of promoter strength across media. Comparison of relative expression of GFP in Y-1190 in **A.** YNBL and supplemented permeate **B**. YNBL and YNBG. Comparison of relative expression of GFP in CBS 6556 and Y-1190 grown in **C**. YNBL at 6 h and 24 h **D**. YNBG at 6 h and 24 h. Each promoter was tested in 4 biological replicates, the IQR method was used to remove outliers and the error bars are reported in standard deviation.

In parallel, we developed three different anhydrotetracycline (aTc) inducible promoters as tools for genetic expression modulation in *K. marxianus*. We introduced two bacterial *tetO* modules into the strongly expressed promoters *TEF3*p, *PGK1*p, and *TDH3*p from *K. marxianus*, and fused those promoters to a mNeonGreen reporter system^34^. This transcriptional unit was used in tandem with a codon-optimized bacterial TetR protein containing a nuclear localization site (NLS) under the control of the strong promoter *SSA3*p. Those constructs were integrated in the genome of *K. marxianus* at the previously published *ARO1* safe harbour site^22^ using the optimized integration protocol presented in the next section, and fluorescence levels upon aTc addition were measured (Fig 4AB). While the p*TEF3*_tetO2_ and p*PGK1*_tetO2_ engineered promoters displayed high induction (19- and 14-fold increase) upon aTc addition, together with tight control in the absence of the inductor, p*TDH3*_tetO2_ showed leaky expression and a lower level of induction (8-fold increase).

**Figure 4.**
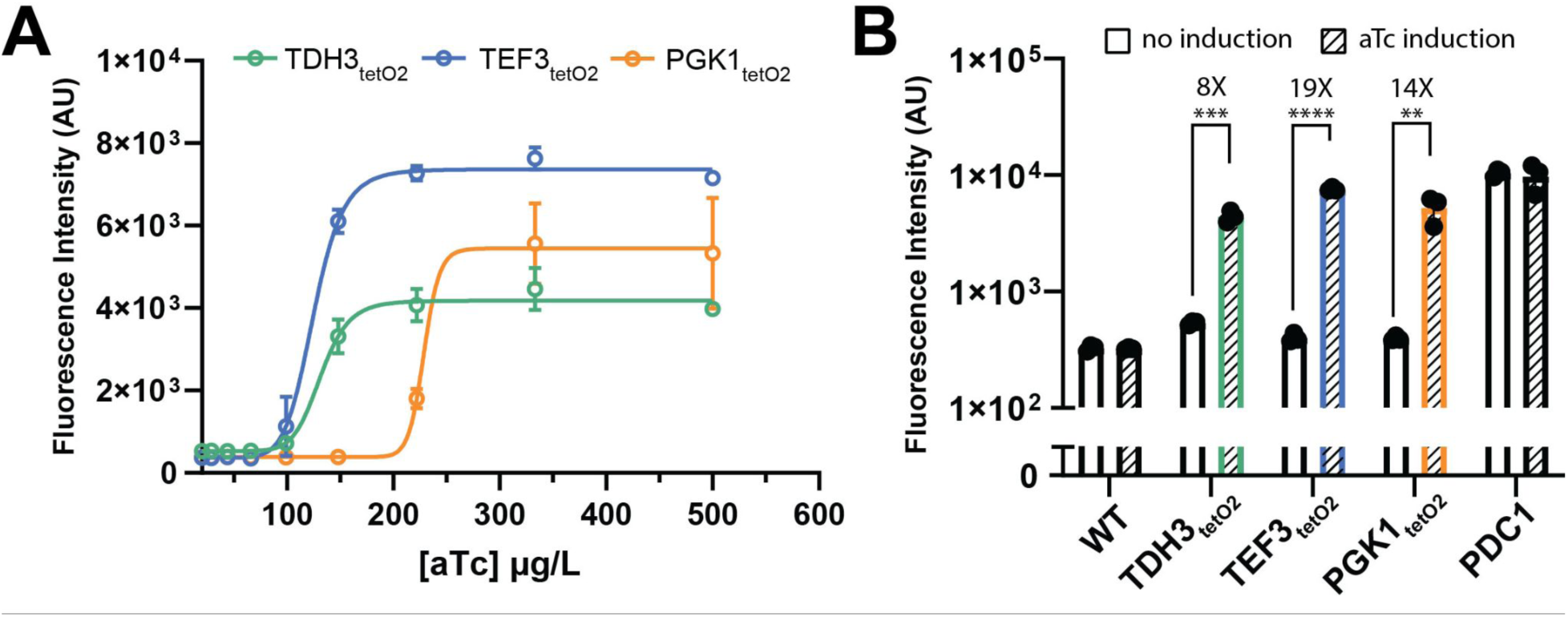
Characterization of aTc-inducible promoters. **A.** Titration of promoter activity upon various aTc concentrations. **B.** Fold change upon maximal induction.

### CRISPR genome editing optimization

Facile genomic engineering of the *K. marxianus* Y-1190 strain is crucial for its use as a platform for biomanufacturing diverse biomolecules. As endogenous gene deletions and genomic integrations are critical for this purpose, we optimized a CRISPR-based integration/deletion protocol for efficient genome editing. We first adapted a GFP-dropout pCas9 plasmid from the *S. cerevisiae* Mylo Toolkit^35^ by swapping the 2µ replication origin with the yeast broad-species pan(OPT)ARS^36^. The resulting plasmid (pMTKM01) was used to build a single guide plasmid targeting the endogenous *URA3* gene from *K. marxianus* Y-1190. We tested whether the transformation of this plasmid was sufficient to integrate a linear donor DNA containing 900 bp of 5’ and 3’ *URA3* homology arms flanking a strongly expressed mNeonGreen. Although this strategy was able to inactivate the *URA3* gene, we were unable to integrate the mNeonGreen cassette (Fig 5A). NHEJ prevalent DNA repair was reported by several studies to interfere with HDR-based genetic integration in different *K. marxianus* strains^22,37,38^. To reduce or inactivate the NHEJ pathway in Y-1190 to favor HDR-based integrations, we generated a NHEJ ligase mutant (ΔDNL4)^39^, and tested the use of a NHEJ chemical inhibitor (NU7026)^40,41^. In parallel, we tested whether prolonged liquid growth with selective pressure for the pCas plasmid after transformation (outgrowth) would favor yeasts with modified Cas9 target sites via HDR, but none of those strategies led to a successful HDR integration or *URA3* inactivation (Fig 5A). We next tested introducing two gRNAs to target different areas of the CDS, a strategy commonly used in mammalian cell culture transformations^42,43^. This double cut approach in combination with the NHEJ inhibition strategies facilitated integrating the mNeonGreen donor DNA at the *URA3* locus (Fig 5A). The most successful protocol, with 100 % of the tested colonies displaying genomic integration, used the two-guide strategy in combination with NHEJ inactivation via genetic deletion, followed by outgrowth. Finally, to verify that the increase in efficiency was due to the use of two guides and was not gRNA sequence specific, we performed the one-guide experiment again with the second gRNA targeting *URA3* (Fig S5). Similarly to the first single gRNA, we were not able to generate any integration, confirming the importance of the two-guide strategy.

**Figure 5.**
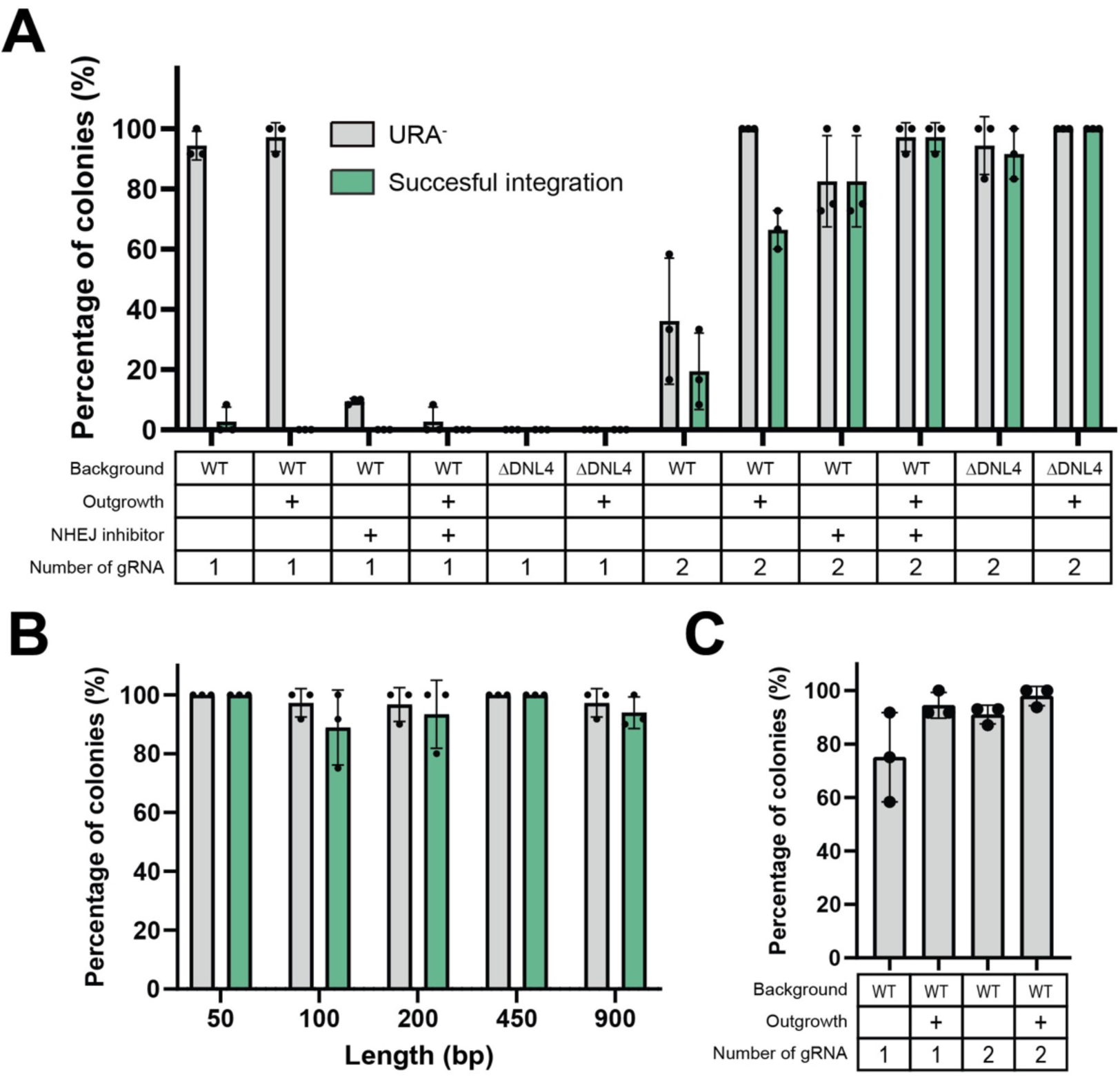
CRISPR-Cas9 genome editing efficiencies in *K. marxianus*. **A.** Optimization of genomic deletion and integration via HDR. **B.** Homologous arm size effect on genomic deletion and integration efficiency via HDR. **C.** Optimization of genomic deletion via NHEJ. Each dot represents a biological replicate and the error bars represent the standard deviation.

We then used our optimized method with outgrowth and *DNL4* deletion to identify the minimal homology arm size necessary for genomic integration. We amplified the mNeonGreen cassette with different sets of primers to generate 900 bp, 450 bp, 200 bp, 100 bp, and 50 bp *URA3* homology arms and tested their efficiencies (Fig 5B). High integration rates were measured for homology arms size as low as 50 bp, consistent with previous reports in other *K. marxianus* strains^25^. Finally, we investigated if gene deletions could be generated only by NHEJ, without providing donor DNA. We tested the effect of the outgrowth and the double cut strategies on this approach. Although a single cut was enough to generate some *URA3* inactivation, 100 % inactivation could be reached either using the double cut or the outgrowth strategies (Fig 5C).

### Testing of diploid stability

Generation of a stable haploid of *K. marxianus* Y-1190 would enable a more streamlined CRISPR protocol as well as the ability to use mating-based assays and tools for genetic manipulation. *K. marxianus* is characterized as homothallic, able to exist as haploid or diploid with the ability to spontaneously switch between mating types^44,45^. This switching occurs via a two-transposase system first identified in *K. lactis* and later confirmed in *K. marxianus*^23,44,46,47^. To stabilize mating competent haploid strains, we used CRISPR-Cas9 to delete the two transposases, *α3* and *Kat1* (Fig S4). We then sporulated the Y-1190ΔKat1Δα3 strain and used two separate methods, classical tetrad dissection and NP-40 based spore retrieval^48^ to generate haploids. Over 100 tetrads were dissected, and hundreds of cells went through the NP-40 based protocol, yet we retrieved only one haploid, JD044, which subsequently autodiplodized (Fig S6). Such autodiploidization has also been noted for *C. albicans* cells after selection for the haploid state^49^. In all media tested diploidized JD044 had a lower growth rate than the wild type Y-1190, thus reducing its utility as a platform for genetic engineering (Fig S7).

### Optimization of *K. marxianus* fermentation in dairy permeate

After identifying a dairy strain of *K. marxianus* highly suitable for genetic manipulation, we evaluated its growth performance in a controlled fermentation environment using small scale bioreactors and dairy permeate. *K. marxianus* Y-1190 cells were first grown on dairy permeate without pH control for 48 hrs to observe the intrinsic biomass yield and pH decline while using this substrate (Fig 6A). At the end of the batch fermentation, strain Y-1190 had a biomass yield of 0.15 g/g lactose, consuming all lactose after around 30 hrs. A previous study from Fonseca *et al.*^50^ showed that the biomass yield of *K. marxianus* in glucose can reach up to 0.51 g/g glucose. Therefore, the biomass yield under our condition was only one-third of the maximum biomass yield in glucose, suggesting limitation of nutrients in the media.

**Figure 6.**
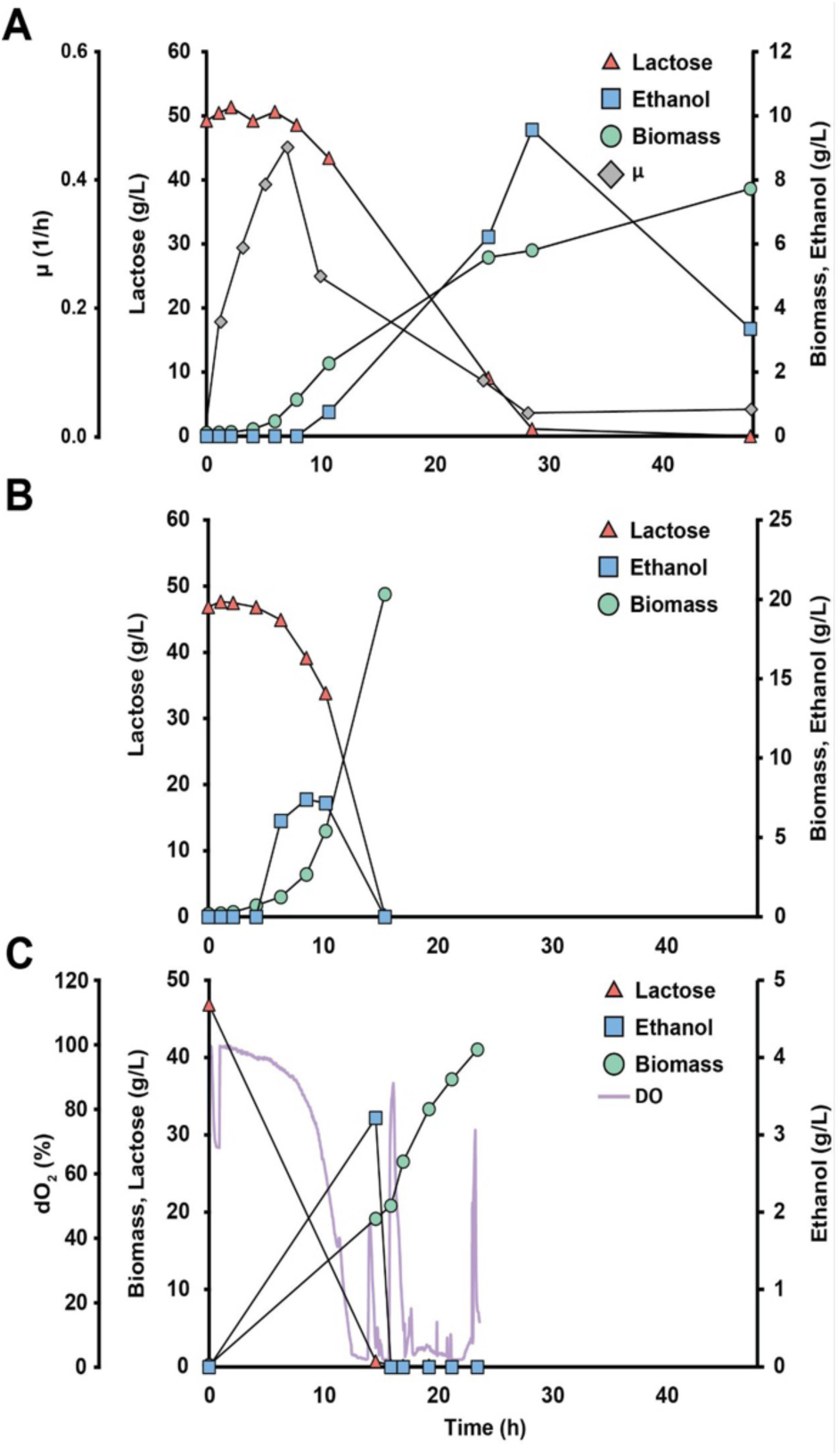
Fermentation profile of *K. marxianus* 1190. **A**. Batch fermentation in non-supplemented dairy permeate. **B.** Batch fermentation in supplemented dairy permeate. **C.** Fed-batch fermentation in supplemented dairy permeate with pH maintained at 4.

To optimize media composition and increase biomass yields we adjusted the composition of the permeate. Based on composition measurement, the total nitrogen from protein in the permeate is only 0.001 g/L (Table 2). The optimum mole of nitrogen per mole of carbon for *K. marxianus* growth is estimated to be 0.07^51^. With around 50 g/L lactose in the permeate (Table 2), the predicted minimum nitrogen needed is 1.69 g/L, indicating a lack of nitrogen in the permeate. Similarly, some minerals and vitamins, such as potassium, phosphorus, sulfur, magnesium, vitamins, and microelements, were also lacking in permeate when compared to the required amounts for the yeast fermentation process^52^. Thus, to increase *K. marxianus* biomass yield, we added into the permeate both urea as a nitrogen source as well as additional nutrients^51^. The batch fermentation of *K. marxianus* Y-1190 using supplemented permeate in a controlled fermentor without pH control showed a maximum uptake rate of lactose of 6.50 g/L lactose/hour (Fig 6B), with no lactose detected after 15 hrs. Under these conditions, a biomass yield of 0.43 g/g lactose was obtained.

Having established the necessary nutrient supplementation for efficient *K. marxianus* fermentation in permeate, we began optimizing the fed-batch phase for biomass yield by balancing the amount of supplied lactose with the oxygen supply of the fermentor. Oxygen limitation was observed during the batch phase, where ethanol was accumulated (Fig 6B). An increase in pH can occur when the rate of ammonium production from urea metabolism is faster than the assimilation of ammonium into biomass^53,54^. This rise in pH has been associated with lactose accumulation ^53,55^, so we adjusted the pH to 4 to optimize the consumption of lactose. Under these conditions we characterized the biomass yield in fed-batch mode using supplemented permeate (Fig 6C). No residual lactose was observed and the biomass yield of *K. marxianus* Y-1190 reached 0.51 g/g lactose, similar to the biomass yield of *K. marxianus* in glucose^50^. This result indicated that a high biomass yield of *K. marxianus* strain Y-1190 can be obtained in supplemented dairy permeate, underlining the potential of this strain being used for the valorization of industrial dairy waste.

## DISCUSSION

There is a growing body of literature for *K. marxianus* that highlights its inherent capacity for industrial fermentation^1^, lignocellulosic biomass degradation^9^, and protein expression^11,56^. New and improved techniques to increase the genetic tractability of this organism have been published^23,25,48,57^, but few studies have harnessed the strain’s inherent ability to metabolize lactose. Although lactose has incredible potential as a feedstock for valorization, few studies have leveraged the ability of *K. marxianus* to metabolize lactose. In Canada alone, dairy companies produce an estimated 30-50 million kilograms of lactose-rich ultrafiltered whey and milk permeate powder annually, while in the U.S., estimates reach 500-600 million kg (Agropur, personal communication, January 2025). Lactose, often considered a low-value byproduct, could play a crucial role in the circular economy - a framework aimed at decoupling economic growth from resource consumption by regenerating and recycling materials^58^. Despite the accessibility of this otherwise low value carbon source, many metabolic engineering studies often focus on yeasts such as *S. cerevisiae*, *Pichia pastoris*, *Pichia occidentalis*, and *Yarrowia lipolytica* that cannot metabolize lactose^6^. Those studies that do use *K. marxianus* often use non-dairy derived strains with less efficient lactose metabolizing properties^9,22,32^. Here we identify *K. marxianus* Y-1190 as having the combined characteristics of robust growth on lactose-containing dairy permeate and tolerance to acid making an excellent basis for an industrially relevant strain (Fig 1).

To begin our domestication process, we sequenced the genome of *K. marxianus* Y-1190. Our genomic sequencing and observation of genome copy number established the diploid nature of this organism (Fig S6). Further, the haplotigs generated by PacBio sequencing point to a diploid organism with considerable heterozygosity. We observed that the primary assembly contained 4,853 predicted genes, while the haplotig assembly contained 4,108 predicted genes, representing 85% of the predicted genes in the primary assembly. This proportion aligns with the total assembly sizes: the primary assembly spans 1.1 × 10⁷ bps, whereas the haplotigs collectively cover 8.2 × 10⁶ bps, representing 85% of the primary assembly’s size across 66 fragments (Table 1). The changes observed at the DNA level do also translate to amino acid changes (Fig S8) suggesting there could be functional differences between the different alleles. The homothallic nature of *K. marxianus* means that it can be *MAT*a, *MAT*alpha, diploid, or a mixed population^6,45^. The preferred ploidy of *K. marxianu*s is thought to be either haploid or diploid dependent on the strain, however, the mechanism of this preference is currently unknown. Previously published work that sequenced *K. marxianus* strain SP1 (NRRL Y50883) using PacBio saw similar high levels of heterozygosity^59^. It is possible that this heterozygosity made generating a stable haploid nearly impossible. Due to the heterologous nature of the genome, the single-nucleotide polymorphisms across alleles may play a role in strain fitness and viability. We were able to obtain one viable clone, JD044, however, we observed that this was not a stable haploid, and though lacking the mating machinery, the genome duplicated to a homozygous diploid state (Fig S6). Losing heterozygosity also cost Y-1190 its high growth rate suggesting that heterozygosity is important for fitness (Fig S7). This was an unexpected outcome as previously published work described Y-1190 as preferring the haploid state, and after removing the mating type-switching transposases, a stable haploid with mating capability was identified^23^. However, ploidy variation in *K. marxianus* has been observed previously. The ploidy of the commonly used strain CBS 6556 has been debated^60^ and short-read sequencing has made it difficult to answer these questions.

**Table 1.** *K. marxianus* Y-1190 summary assembly. Statistics and annotation of primary assembly and haplotigs.

|  | Primary Contigs | Haplotigs |
| --- | --- | --- |
| Sequences | 10 | 66 |
| GC% | 40.1 % | 40.6 % |
| N50 | 1,590,052 | 935,761 |
| Assembly Length | 10,951,246 | 9,309,754 |
| Gene Annotations | 4853 | 4108 |

To enhance metabolic engineering tools in *K. marxianus*, we characterized 23 promoters from strains Y-1190 and CBS 6556 across three media types: supplemented milk permeate, YNBL, and YNBG. Twenty-one of the 23 promoters were sourced from Lang *et al.*^20^ allowing a comparison of promoter strength across strains of *K. marxianus* (Fig 8A). Notably, three high-expressing promoters (*pNC1*, *pTEF3*, *pPGK1*), one mid-expressing promoter (*pGLK1B*), and one low-expressing promoter (*pRPE1*) exhibited consistent rankings (Fig 7). Overall, the Pearson’s correlation coefficients between our findings and Lang *et al.* were weakly positive.

**Fig 7.**
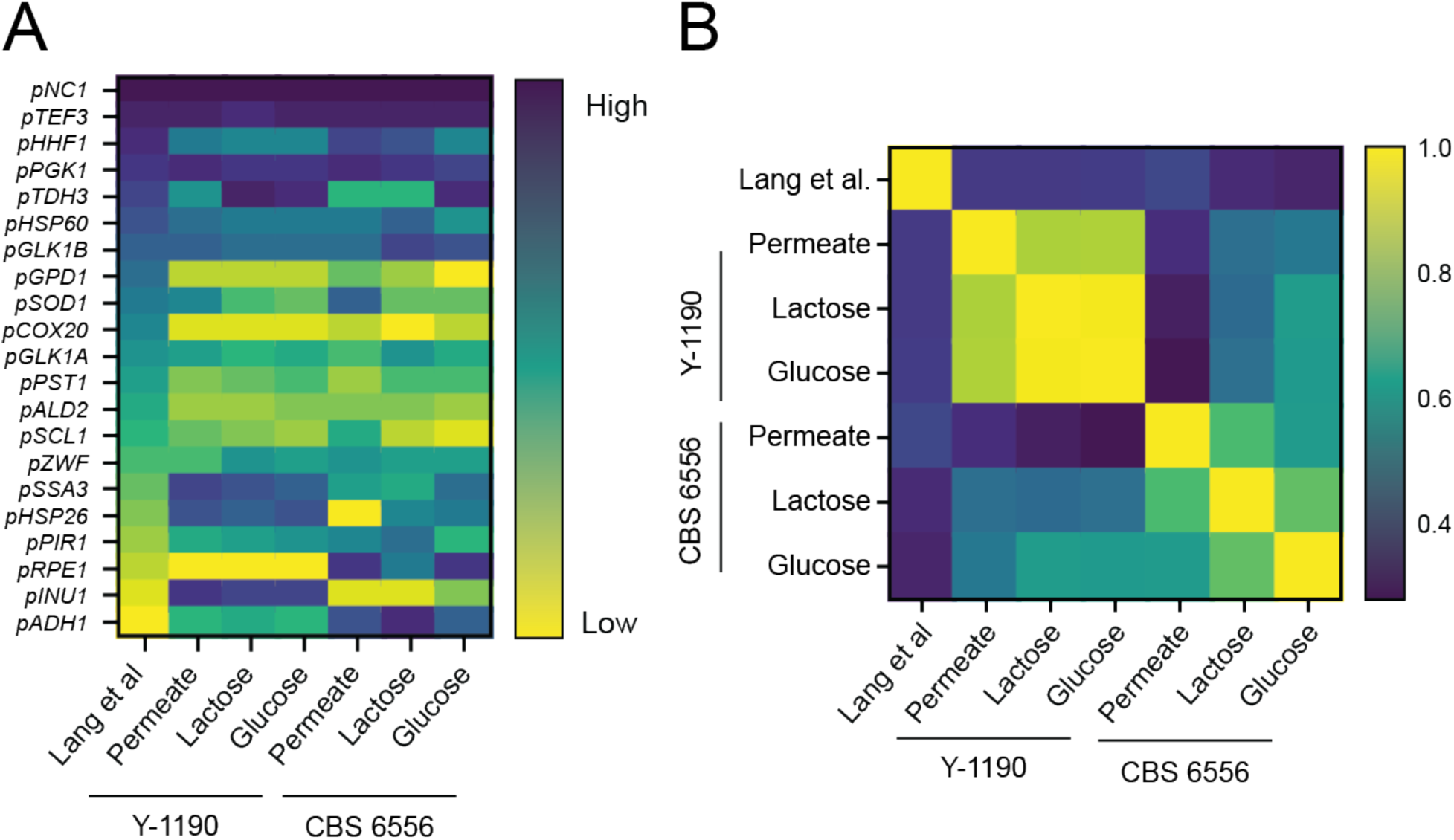
Promoter comparison to Lang *et al*. 2020. **A.** Common promoters used in both studies were ranked, with dark blue representing the highest expression and yellow representing the lowest. **B.** The ranked lists were compared by calculating Pearson’s correlation coefficient, shown in a heatmap. The correlation values range from *r* = 1.0 (yellow) to *r* = 0.4 (dark blue).

Rajkumar *et al.* tested 16 promoters from CBS 6556, sharing only five with our study, complicating direct comparisons. Our study also overlaps with only four promoters from the *S. cerevisiae* toolkit, yet *pTDH3* and our highest expressor, *pNC1* (homologous to *S. cerevisiae pCCW12*), are noteworthy for maintaining high expression both in *S. cerevisiae* and *K. marxianus*^33^.

Promoter strength comparisons within this study reveal greater similarity across strains than media types (Fig 8B). For example, Y-1190 promoter strengths in supplemented permeate and YNBG were more similar than between Y-1190 and CBS 6556 in YNBG, suggesting strain-specific regulatory differences. Variations in transcription factors or regulatory proteins could explain these discrepancies, as Y-1190 is diploid while CBS 6556 is haploid, potentially affecting expression through gene dosage or chromatin structure^61,62^. This is reinforced by a previous study that showed promoters maintain their relative activity across different media types^63^.

*pINU1* was identified as a differentially expressed promoter; although typically low in the absence of inulin, it is highly expressed in Y-1190 (Fig 3, Fig 7A). This suggests regulatory differences or sequence variations affecting activator binding. Overall, the relationship between promoter expression in CBS 6556 and Y-1190 appears linear across media, except in supplemented permeate due to CBS 6556’s poor growth (Fig 7B, Fig S4).

*K. marxianus*, like many non-conventional yeasts, is known to prefer non-homologous end joining (NHEJ) to homology-directed repair (HDR) for DNA repair mechanisms^22,37,38^. Through NHEJ, CRISPR can be used to make insertions or deletions at the gRNA target site that can disrupt the gene but cannot be used for integrations of exogenous DNA (Fig 5). This makes confirmation of genetic disruptions both time consuming and expensive, as screening can only be done by phenotype or sequencing. Many groups have worked to make CRISPR-Cas9 more effective in *K. marxianus* through deleting the NHEJ machinery^22,23,37^. We have combined that technique with the addition of an NHEJ inhibitor and a double guide method commonly used in mammalian cell CRISPR studies^40–43^. This combination of techniques directed 100 % replacement of km*URA3* with mNeonGreen using homology arms of only 50 bp (Fig 5). These results, in combination with our sequencing data, promoter library characterization, inducible promoters, and fermentation optimization have expanded the genetic toolkit for *K. marxianus*, increasing the potential for metabolic engineering applications and enabling more precise control over gene expression for improved biotechnological outcomes.

## MATERIALS AND METHODS

### *Kluyveromyces marxianus* strain selection and media optimization

Nineteen strains of *K. marxianus*, and one control strain of *Saccharomyces cerevisiae* (CEN.PK), were revived from frozen stocks and cultured on YPD (10 g/L Bacto Yeast Extract, 20 g/L Bacto peptone, and 20 g/L glucose) agar plates at 30 °C. Three independent colonies of each strain were selected to inoculate 800 μl of YPD media in a 96-well deep plate. Once the cultures reached visible turbidity, all the strains were back-diluted into 800 μl of YPD in a 96-well deep plate and incubated for 8 hrs to reach an OD_600_ of 1.8. After an 8-hr incubation, a diluted intermediate plate was created to achieve an OD_600_ of 0.9. A 10 μL inoculum from the diluted plate was used to inoculate 170 μl of the relevant growth medium. The final volume of each well was 180 μL, and the initial OD_600_ of all strains was 0.05. Growth rates (μ_max_) were calculated in R using the growth rates package^64^.

To test the growth of *K. marxianus* strains on various sugars, strains were grown in YNB-Glucose (YNBG, 20 g/L), YNB-Lactose (YNBL, 20 g/L), and in supplemented dairy permeate, a filtered permeate from industrial dairy processing waste (Agropur Cooperative, Table 2) supplemented with vitamins and microelements (8.36 g/L urea, 2.07 g/L KH_2_PO_4_, 1.15 g/L MgSO_4_-7H_2_O, 1 g/L NiSO_4_-6H_2_O, 5 mL/L vitamin stock, and 5 mL/L microelements). The composition of the vitamin and microelement stock was described previously^65^. For the acid tolerance test, 5 g/L of fumaric acid was added to the supplemented permeate, and 12 N HCL was added dropwise to the final pH of 2.2.

**Table 2.**
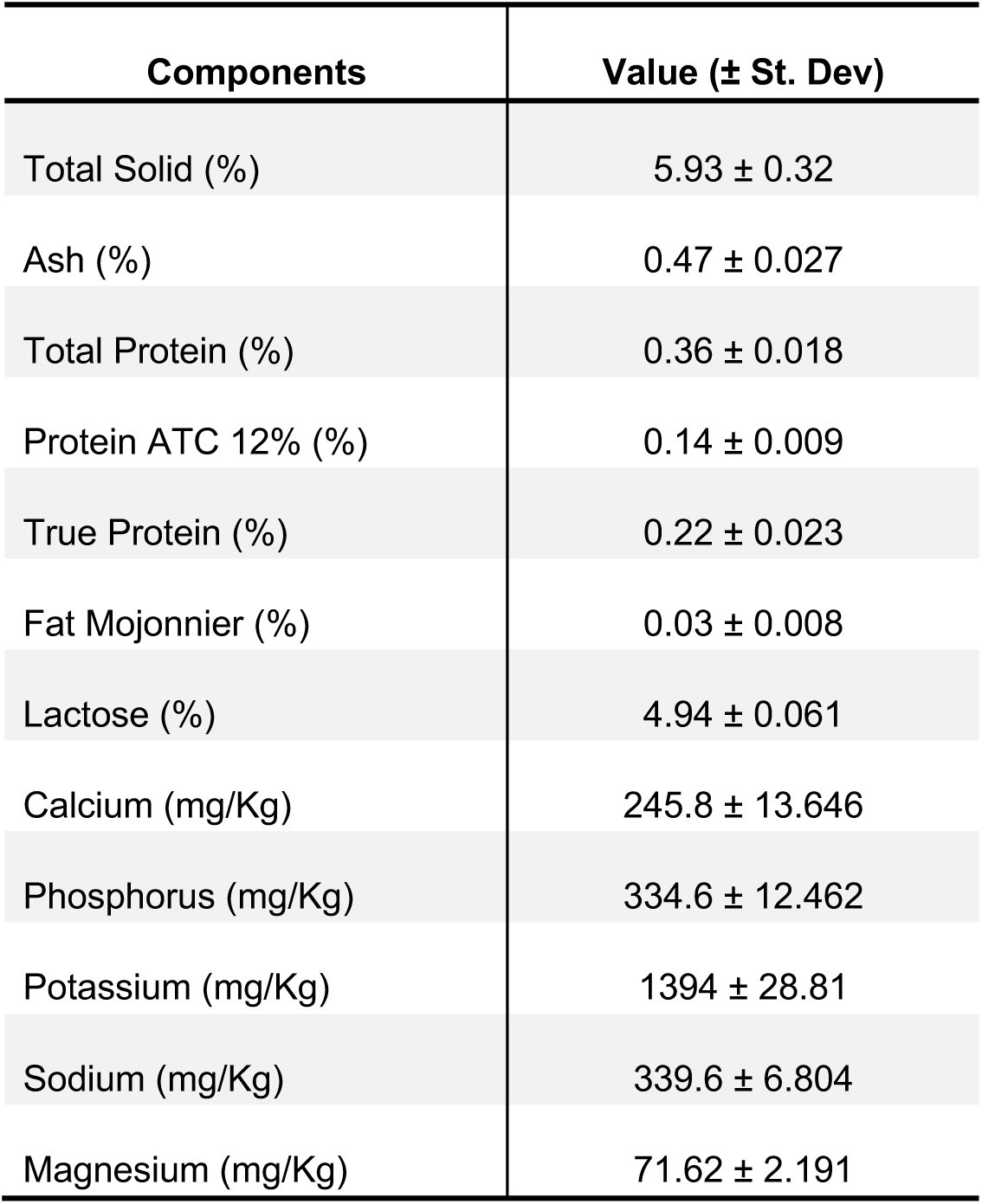
Dairy permeate dry composition analyzed by Inductively Coupled Plasma Spectroscopy conducted by Mérieux NutriSciences, Dorval, Quebec.

| Components | Value (± St. Dev) |
| --- | --- |
| Total Solid (%) | 5.93 ± 0.32 |
| Ash (%) | 0.47 ± 0.027 |
| Total Protein (%) | 0.36 ± 0.018 |
| Protein ATC 12% (%) | 0.14 ± 0.009 |
| True Protein (%) | 0.22 ± 0.023 |
| Fat Mojonnier (%) | 0.03 ± 0.008 |
| Lactose (%) | 4.94 ± 0.061 |
| Calcium (mg/Kg) | 245.8 ± 13.646 |
| Phosphorus (mg/Kg) | 334.6 ± 12.462 |
| Potassium (mg/Kg) | 1394 ± 28.81 |
| Sodium (mg/Kg) | 339.6 ± 6.804 |
| Magnesium (mg/Kg) | 71.62 ± 2.191 |

### Transformation

*K. marxianus* was transformed using the Gietz method with changes to heat shock temperature and time^30^. Briefly, cells from 3 OD_600_ units of log phase cultures were harvested by centrifugation (1860 g) for 3 min, washed with dH_2_O, and transformed in a total volume of 150 μl using the concentrations as described previously^30^. The transformations were incubated at room temperature for 20 min prior to heat shock at 47 ℃ for 15 min. Transformations were recovered overnight in YPD at 30 ℃ without shaking and subsequently back diluted 1:5 into YPD and the appropriate antibiotic for 24-48 h for optional outgrowth, before plating on selection media.

### Genome sequencing and bioinformatics

#### Yeast strains and library preparation

*K. marxianus* strain Y-1190 was subjected to whole genome sequencing. Y-1190 was cultured overnight in 5 mL YPD at 30 ℃ and 300 rpm. Cells were harvested by centrifugation and DNA was purified using the MasterPure Yeast DNA Purification Kit (Lucigen) following the manufacturer’s instructions for cells harvested from liquid cultures except that DNA was eluted in nuclease-free H_2_O rather than TE buffer to avoid EDTA contamination. DNA quantity was measured using a Qubit 4 fluorometric assay (Invitrogen) and was 560 ng/μL. All yeast strains used in this study can be found in Table S3.

#### Long-read sequencing, genome assembly and annotation

Genomic DNA was sequenced (HistoGenetics Inc., Ossining NY) using the PacBio SMRT Sequel II platform following library prep using the SMRTbell HiFi kit (Pacific Biosciences) and generated 7.31 Gb reads for *K. marxianus* Y-1190. Demultiplexing and *de novo* genome assembly was achieved using the SMRTlink *de novo* assembly pipeline^66^.

Low complexity and highly repetitive regions were masked using RepeatMasker^67^. AUGUSTUS v3.4.0^68^ was used for *ab initio* gene prediction. To assign identity, predicted genes were queried by BLASTX 2.10.0+ or DIAMOND ^69^ against the non-redundant database v5 or v2023-09-21, respectively, using default parameters in Omicsbox with the taxonomy filter set to 4910 (*Kluyveromyces*). Predicted genes were aligned with BLAST against Saccharomycetalae to assign identity, annotation performed using InterProScan along with GO mapping and annotation. Predicted proteins were queried against the AybraH yeast protein database however, since *K. marxianus* isn’t present in this database, NCBI BLAST/diamond returned better matches. Annotated genome files are available at NCBI (BioProject: PRJNA957743, Accession: JBJBSF000000000). BUSCO (v5.4.5) completeness scores were used to assess genome completeness using the following parameters: Lineage Saccharomycetes, Mode Transcriptome, and e-value < 10×10^-3^ along with the OrthoDB v10 database.

### Promoter set characterization

#### Promoter selection

Previous toolkits make use of sets of promoters of varying strength to provide tunable gene expression – e.g. the Yeast Toolkit^33^ provides 23 swappable promoters. We selected 23 *K. marxianus* promoters with a wide range of strengths based on previous reports^20,22^.

Two *K. marxianus* promoter parts from Rajkumar *et al.* were obtained from Addgene ^22^. Twenty-one promoters from Lang *et al*. were a generous gift from Dr. Ian Wheeldon (University of California Riverside). These promoters were not in a modular cloning format and were cloned into the YTK MoClo format^33^, including removing unwanted type IIS restriction sites, before building and testing (pMTKM09-pMTKM29). All primers used in this study can be found in Table S4.

The three anhydrotetracycline (aTc) inducible promoters were designed by replacing endogenous sequences with two bacterial *tetO* modules (TCCCTATCAGTGATAGAGA) separated by two nucleotides (AT) 10 nt after the identified TATA box of the strongly expressed promoters *TEF3*p, *PGK1*p, and *TDH3*p from *K. marxianus.* Those engineered promoters were amplified from *K. marxianus* Y-1190 genomic DNA and any BsaI or Esp3I sites present within their sequences were removed using truncated primers. Esp3I sequences were added with primer overhangs at 3’ and 5’ ends and the resulting amplicons were assembled with GoldenGate into level 0 plasmids from the Modular cloning suite (pMTKM30-pMTKM32) ^33^. The engineered promoters were next fused to mNeonGreen and the *K. marxianus PDC1*t terminator following the MoClo syntax. In parallel, a *K. lactis* codon-optimized version of the bacterial repressor gene *tetR* fused to the Nuclear Localization Signal SV40 was ordered as a synthetic gene (Twist Bioscience) and cloned into a level 0 MoClo entry plasmid. The repressor was next fused to the *SSA3*p promoter and the *INU1*t terminator from *K. marxianus* following the MoClo syntax. Finally, the two transcriptional units were fused together within a plasmid containing the 5’ and 3’ homology arms for integration into the *ARO1* safe harbour site, as previously described^22^. Those plasmids were digested with NotI-HF and used as donor DNA together with a pCas plasmid containing two guides targeting the *ARO1* locus for genomic integration (see section genome engineering for details). Finally, the resulting strains were cured of their pCas plasmids and inoculated overnight in YPD prior to their dilution at an OD_600_ of 0.1 in Synthetic Complete (SC) medium. Cells were grown in deep-well plates for 6 hrs until OD_600_ of ∼1 and fluorescence was measured using a plate reader (Infinite200, Tecan).

#### Promoter reporter plasmid construction

Promoter activity reporter plasmids consisted of the promoters listed in Table S2, 5’ to the sf*GFP* from the Yeast ToolKit paper, and the Km*PGK*t terminator from the *K. marxianus* biological parts paper^22,33^. These plasmids were created using Golden Gate^70^. Briefly, 40 fmol of each insert was combined with 20 fmol of backbone plasmid (YTK083) along with 0.5 μL BsaI-HF (NEB), 0.5 μL T4 ligase, 1.5 uL 10x T4 ligase buffer, and water to a 15 μL total volume. Assembly was carried out by digestion at 37 °C for 5 min followed by ligation at 16 °C for 5 min for 30 cycles. A final 37 °C incubation was done for 60 min digestion followed by an inactivation step of 80 °C for 15 min. The resulting plasmids were transformed into chemically competent *E. coli* DH5alpha cells and stored in glycerol at -80 °C for later use. Plasmids were sequenced by nanopore sequencing (Plasmidsaurus). Once confirmed by sequencing these 23 plasmids were transformed into *K. marxianus* strains Y-1190 and CBS 6556 for promoter strength testing. All plasmids used and created in this study can be found in Table S5.

#### Promoter strength testing

Four independent colonies from each promoter plasmid were selected and grown overnight in 700 μl of YPD with 100 μg/mL nourseothricin selection. Cells from overnight cultures were washed twice in water and an intermediate 96-well plate was created at an OD_600_ of 1 in water. This plate was used to seed our experiment in 96 deep well plates at an OD_600_ of 0.1 in either YNBL, YNBG, and 0.5x supplemented dairy permeate. These plates were grown at 30 ℃ with shaking at 300 rpm for 6 and 24 hrs at which point fluorescence and OD_600_ were measured using a plate reader (Infinite200, Tecan). The interquartile range (IQR) was calculated for each promoter and data points that fell outside 1.5*IQR on either side were removed. Figures were created in R.

### Genome editing using CRISPR-Cas9

A series of SpCas9/gRNA vectors originally published by Ryan *et al*., 2014^71^, and later simplified by Bean 2022^35^, were modified for optimal functionality in *K. marxianus* by replacing the 2μ sequence with a panARS(opt) replication origin^36^. The resulting plasmids display a BsaI GFP-dropout for easy gRNA cloning via Golden-Gate and are available on Addgene (pMTKM01-pMTKM08, see Table S5). To design the different gRNAs used in this study, we used CCTop^72^, and selected guides displaying at least a CRISPRater score^73^ of 0.60. Golden-Gate cloning was performed as previously described to introduce one guide into the vector^35^. To express two gRNA from the same transcriptional unit within one vector, we used the tRNAgly self-splicing module as previously reported^74^. We used the already published pCD4 plasmid^75^ containing a tRNAgly together with a SpCas9 sgRNA scaffold as a template for amplification with primers containing the two sgRNAs followed by two BsaI sites as overhangs. The amplicon generated was next purified and 80 fmol was added to a Golden-Gate reaction containing 20 fmol of vector as described in the previous section.

The donor DNA used for the optimization of the protocol for CRISPR-Cas9 genome editing with HDR was amplified by PCR from the pMT187 and pMT188 plasmids. Briefly, *PDC1*p, mNeonGreen, *PDC*t, and the 3’ and 5’ regions of the *URA3* genes were amplified by PCR with overlapping overhangs and cloned into the pJET plasmid (Thermo Fisher Scientific) according to the manufacturer’s instructions. The plasmid was next used as a template to generate donor DNA with different recombination arm sizes with different sets of primers. The ΔDNL4 strain was generated following a protocol previously published^22^.

For the deletion/integration test of the *URA3* locus, *K. marxianus* was transformed as described in the previous section with 500 ng of Cas9 plasmid vector. For integrations, 500 ng of Cas9 plasmid vector and 1-2 μg of donor DNA were provided. After recovery overnight in 500 µl of YPD, cells were plated on YPD with selection for the Cas9 plasmid. For outgrowth, 100 µl of the recovery culture was inoculated into 400 µl of YPD with selection for the Cas9 plasmid and grown for 72 hrs in deep-well plates. For NHEJ chemical inhibition, NU7026 (Cayman Chemicals) was added at a final concentration of 25 µg/ml at every stage of the transformation until plating. After 48 hrs of growth on selective media at 30 °C, at least 12 colonies per technical replicate were picked with a colony picker (Qpix400, MolecularDevice) and inoculated in 96-well microplates in SC medium supplemented or not with 5-FOA at a final concentration of 1 g/L. Cells were grown at 200 rpm for 16 hrs at 30 °C, and OD_600_ was measured together with green fluorescence to evaluate *URA3* inactivation and mNeonGreen integration, respectively.

### Investigation of strain haploidization

To test the diploid-haploid transition in *K. marxianus* strain Y-1190, cells were sporulated as described by Wu et. al.^48^, with minor modifications. An overnight culture grown at 30 °C in YPD was used to inoculate 5 ml of YPA (20 g/L potassium acetate, 20 g/L bactopeptone, 10 g/L yeast extract) at an OD_600_ of 0.2. The cells were grown for 3-5 hrs to a final OD_600_ of 0.8-1.0, washed once with sterile dH_2_O (5,000 x g, 5 min), and suspended in 5 ml of 2 % potassium acetate. Following 16 hrs of incubation at 30 °C with shaking, approximately 50 % of the cells had formed asci. Cells from 1 ml of the sporulated culture were washed twice with dH_2_O and suspended in a final volume of 500 μl. To eliminate non-sporulated cells, cell walls were digested by the addition of 25 μl of 5 unit/μl 100-T Zymolyase (Zymo Research, E1004) with 5 μl of β-mercaptoethanol. The mixture was incubated at 30 °C for 30 min without shaking followed by the addition of 200 μl of 1.5 % NP-40 and incubation at 30 °C for 30 min. Asci were dispersed by incubation in a sonicating water bath for 30 sec intervals (10-15 repetitions). Once sufficient dispersal was achieved, the mixture was washed twice with dH_2_O, serially diluted to 10^-5^, and 100 μl was plated on YPD. Colonies were then screened by colony-PCR both to identify potential haploids and to determine their mating type.

To trap potential haploids in a mating competent form, the transposases *α3* and *Kat1* were targeted by CRISPR-NHEJ using the two-gRNA strategy described above. Guides were designed such that successful inactivation caused the removal of a 1532 bp (*Kat1*) or 652 bp (*α3*) sequence, resulting in a size difference that can be detected by gel electrophoresis after PCR amplification of the gene region. Strain Y-1190 was transformed with the construct targeting *Kat1* (pLN026), followed by plasmid curing and subsequent transformation with the construct targeting *α3* (pLN028). The resulting stain (JD032) had a truncation in both *α3* and *Kat1* as verified by PCR and by sequencing.

Propidium iodide staining followed by flow cytometry^76^ was used to determine the ploidy of potential haploid strains recovered from sporulation. Samples were analyzed using an Accuri C6 flow cytometer with autosampler. Known *S. cerevisiae* diploid and haploid strains, as well as the *K. marxianus* Y-1190 parental diploid strain were used as controls for comparison with candidate haploids.

### Cultivation of *K. marxianus* in bioreactor

Controlled fermentations were conducted in 3 L BioBundle bioreactors (Applikon). The fermentation temperature was maintained at 30 °C. Batch fermentations were performed without controlling the pH, while fed-batch fermentation pH was controlled at pH 4.0 by automated titration with 4 N NaOH. Dissolved oxygen was maintained at 35 % air saturation with an aeration rate of 1 L/min. Concentrations of O_2_ and CO_2_ in the off-gas were analyzed using a Tandem Multiplex Gas Analyzer (Magellan BioTech). Bioreactor inoculum was grown in a 500 mL shake flask containing 50 mL of batch medium at 30 °C overnight. Cells were washed and suspended in 0.9% NaCl before inoculation in 1 L of batch medium with starting OD_600_ of ∼0.3. As a batch medium, filtered dairy permeate with or without supplementation was used as described above. For fed-batch fermentation, the bioreactor was operated in batch mode until lactose exhaustion, followed by the fed-batch phase with the automated addition of a feeding medium. For the feeding medium, three times ultrafiltration-concentrated permeate (Agropur Cooperative) with supplementation (23.07 g urea, 6.19 g KH_2_PO_4_, 3.45 g MgSO_4_·7H_2_O, 3 g NiSO_4_·6H_2_O, 22.5 mL vitamin stock, and 22.5 mL trace element stock per liter) was used. Nickel was added to promote urea utilization as mentioned by Dubencovs et.al^51^. Samples for OD measurements and HPLC analysis were collected every hour for eight hours per day with the two-day total fed-batch running time. Dry cell weight (g/L) was calculated using a gravimetrically determined conversion factor of 0.22 g/L per OD_600_. Bioreactor data were collected and analyzed using BioXpert.V2 (Applikon).

Lactose, glycerol, and ethanol concentrations were analyzed from 10 µL of fermentation samples diluted 1:10 in dH_2_O using HPLC (Agilent 1290 Infinity II) equipped with an Aminex Fast Acid Analysis column (100 ⨯ 7.8 mm, Bio-Rad) and a 10 mM H_2_SO_4_ mobile phase at a flow rate of 0.8 mL/min. Lactose, glycerol, and ethanol were detected using RI, while fumaric acid was detected using a UV detector at 210 nm. Chromatograms were collected and analyzed using OpenLab CDS version 2.7 (Agilent Technologies).

## Supporting information

Thornbury et al Supplementary Tables

Thornbury et al Supplementary Figures

## ACKNOWLEDGMENTS

We thank Dr Ian Wheeldon for the generous gift of his *K. marxianus* promoter library. We thank Lallemand for the generous gift of *K. marxianus* strains L381 and L388. This work was financially supported by Genome Canada, Genome Québec and Agropur. M.T. was supported by an NSERC Vanier Canada Graduate Scholarship, A.K. was supported by a Concordia Horizon Postdoctoral Fellowship, S.J. was supported by an NSERC Canada Graduate Scholarship, J.C.U. was supported by a Lallemand Postdoctoral Fellowship in Bioprocessing. M.W was supported by an NSERC Canada Research Chair, and V.J.J.M. was supported by a Concordia University Research Chair.

## AUTHOR CONTRIBUTIONS

Conceptualization of the project was carried out by VJJM, MW, and MP, while formal analysis was performed by MT, AK, ISM, SJ, JCU, JD, and JJ. Funding acquisition was managed by VJJM, MW, and MP, and investigation was conducted by MT, AK, ISM, SJ, JCU, JD, JJ, LN, and GR. The project administration was facilitated by VJJM, MW, MT, and AK. Software was organized and written by ISM, JCU, and SJ, while resources were provided by VJJM, MW, and MP. Supervision was handled by VJJM and MW. Validation was conducted by MT, AK, SJ, and JCU, and visualization was done by MT, AK, ISM, SJ, and JCU. The original draft of the writing was produced by MT, AK, JCU, ISM, SJ, and JD, with the review and editing process carried out by VJJM, MW, and MT.

## ADDITIONAL INFORMATION

Supplementary information is available in the online version of the paper. Correspondence and requests for materials should be addressed to V.J.J.M.

## DATA AVAILABILITY

The data are available from the corresponding author upon reasonable request.

## REFERENCES

1. Bilal, M. et al. Bioprospecting *Kluyveromyces marxianus* as a robust host for industrial biotechnology. Front. Bioeng. Biotechnol. 0, 562 (2022).

2. Groeneveld, P., Stouthamer, A. H., Westerhoff, H. V & Westerhoff, H. V. Super life-how and why ‘cell selection’ leads to the fastest-growing eukaryote. Authors J. Compil. ^a^ 276, 254–270 (2009).

3. Chang, J. J. et al. A thermo- and toxin-tolerant kefir yeast for biorefinery and biofuel production. Appl. Energy 132, 465–474 (2014).

4. Anderson, P. J., McNeil, K. & Watson, K. High-efficiency carbohydrate fermentation to ethanol at temperatures above 40°C by *Kuyveromyces marxianus* var. *marxianus* isolated from sugar mills. Appl. Environ. Microbiol. 51, 1314–1320 (1986).

5. Nonklang, S. et al. High-temperature ethanol fermentation and transformation with linear DNA in the thermotolerant yeast *Kluyveromyces marxianus* DMKU3-1042. Appl. Environ. Microbiol. 74, 7514 (2008).

6. Kurtzman, C. P., Fell, J. W. & Boekhout, T. The Yeasts, a Taxonomy Study. (Elsevier Science, 2011).

7. Ha-Tran, D. M. et al. Construction of engineered RuBisCO *Kluyveromyces marxianus* for a dual microbial bioethanol production system. PLoS One 16, e0247135 (2021).

8. Christensen, A. D., Kádár, Z., Oleskowicz-Popiel, P. & Thomsen, M. H. Production of bioethanol from organic whey using *Kluyveromyces marxianus*. J. Ind. Microbiol. Biotechnol. 38, 283–289 (2011).

9. Goshima, T. et al. Bioethanol production from lignocellulosic biomass by a novel *Kluyveromyces marxianus* strain. Biosci. Biotechnol. Biochem. 77, 1505–1510 (2013).

10. Hensing, M., Vrouwenvelder, H., Hellinga, C., Baartmans, R. & van Dijken, H. Production of extracellular inulinase in high-cell-density fed-batch cultures of *Kluyveromyces marxianus*. Appl. Microbiol. Biotechnol. 42, 516–521 (1994).

11. Zhang, Y. et al. Inulinase hyperproduction by *Kluyveromyces marxianus* through codon optimization, selection of the promoter, and high-cell-density fermentation for efficient inulin hydrolysis. Ann. Microbiol. 69, 647–657 (2019).

12. Zhou, H. X., Xin, F. H., Chi, Z., Liu, G. L. & Chi, Z. M. Inulinase production by the yeast *Kluyveromyces marxianus* with the disrupted MIG1 gene and the over-expressed inulinase gene. Process Biochem. 49, 1867–1874 (2014).

13. Aggelopoulos, T. et al. Solid state fermentation of food waste mixtures for single cell protein, aroma volatiles and fat production. Food Chem. 145, 710–716 (2014).

14. Alasmar, R. et al. Isolation of a novel *Kluyveromyces marxianus* strain QKM-4 and evidence of its volatilome production and binding potentialities in the biocontrol of toxigenic fungi and their mycotoxins. ACS Omega 5, 17637–17645 (2020).

15. Inokuma, K. et al. Complete genome sequence of *Kluyveromyces marxianus* NBRC1777, a nonconventional thermotolerant yeast. Genome Announc. 3, (2015).

16. Limtong, S., Sringiew, C. & Yongmanitchai, W. Production of fuel ethanol at high temperature from sugar cane juice by a newly isolated *Kluyveromyces marxianus*. Bioresour. Technol. 98, 3367–3374 (2007).

17. Teofilo, H., Miguel, U. & Irma, F. Descripción de una especie nueva de *Hansenula Y* una variedad nueva de *Candida parapsilosis* aisladas del pozol. Bol. Soc. Mex. Mic 7, (1973).

18. Ortiz-Merino, R. A. et al. Ploidy variation in *Kluyveromyces marxianus* separates dairy and non-dairy isolates. Front. Genet. 9, (2018).

19. Market Research Future. Whey Permeate Market Size Worth USD 970.6 Million by 2028. https://www.globenewswire.com/en/news-release/2022/05/04/2435465/0/en/Whey-Permeate-Market-Size-Worth-USD-970-6-Million-by-2028-Witnessing-a-CAGR-of-4-05-Report-by-Market-Research-Future-MRFR.html (2022).

20. Lang, X., Besada-Lombana, P. B., Li, M., Da Silva, N. A. & Wheeldon, I. Developing a broad-range promoter set for metabolic engineering in the thermotolerant yeast *Kluyveromyces marxianus*. Metab. Eng. Commun. 11, (2020).

21. Kumar, P., Sahoo, D. K. & Sharma, D. The identification of novel promoters and terminators for protein expression and metabolic engineering applications in *Kluyveromyces marxianus*. Metab. Eng. Commun. 12, e00160 (2021).

22. Rajkumar, A. S., Varela, J. A., Juergens, H., Daran, J.-M. G. & Morrissey, J. P. Biological parts for *Kluyveromyces marxianus* synthetic biology. Front. Bioeng. Biotechnol. 7, 97 (2019).

23. Cernak, P. et al. Engineering *Kluyveromyces marxianus* as a robust synthetic biology platform host. MBio 9, (2018).

24. Juergens, H. et al. Genome editing in *Kluyveromyces* and *Ogataea* yeasts using a broad-host-range Cas9/gRNA co-expression plasmid. FEMS Yeast Res. 18, (2018).

25. Rajkumar, A. S. & Morrissey, J. P. Protocols for marker-free gene knock-out and knock-down in *Kluyveromyces marxianus* using CRISPR/Cas9. FEMS Yeast Res. 22, 67 (2022).

26. Varela, J. A. et al. Origin of lactose fermentation in *Kluyveromyces lactis* by interspecies transfer of a neo-functionalized gene cluster during domestication. Curr. Biol. 29, 4284–4290.e2 (2019).

27. Tran, V. G. et al. An end-to-end pipeline for succinic acid production at an industrially relevant scale using *Issatchenkia orientalis*. Nat. Commun. 2023 141 14, 1–14 (2023).

28. Werpy, T. & Petersen, G. Top Value Added Chemicals from Biomass Volume I. Us Nrel http://www.osti.gov/scitech//servlets/purl/15008859-s6ri0N/native/ (2004) doi:10.2172/15008859.

29. Sauer, M., Porro, D., Mattanovich, D. & Branduardi, P. Microbial production of organic acids: expanding the markets. Trends Biotechnol. 26, 100–108 (2008).

30. Gietz, R. D. & Schiestl, R. H. High-efficiency yeast transformation using the LiAc/SS carrier DNA/PEG method. Nat. Protoc. 2007 21 2, 31–34 (2007).

31. Kuznetsov, D. et al. OrthoDB v11: annotation of orthologs in the widest sampling of organismal diversity. Nucleic Acids Res. 51, D445–D451 (2023).

32. Varela, J. A. et al. Polymorphisms in the LAC12 gene explain lactose utilisation variability in *Kluyveromyces marxianus* strains. FEMS Yeast Res. 17, (2017).

33. Lee, M. E., DeLoache, W. C., Cervantes, B. & Dueber, J. E. A Highly characterized yeast toolkit for modular, multipart assembly. ACS Synth. Biol. 4, 975–986 (2015).

34. Shaner, N. C. et al. A bright monomeric green fluorescent protein derived from *Branchiostoma lanceolatum*. Nat. Methods 10, 407–409 (2013).

35. Bean, B. D. M., Whiteway, M. & Martin, V. J. J. The MyLO CRISPR-Cas9 toolkit: a markerless yeast localization and overexpression CRISPR-Cas9 toolkit. G3 Genes|Genomes|Genetics 12, jkac154 (2022).

36. Liachko, I. & Dunham, M. J. An autonomously replicating sequence for use in a wide range of budding yeasts. FEMS Yeast Res. 14, 364–367 (2014).

37. Nambu-Nishida, Y., Nishida, K., Hasunuma, T. & Kondo, A. Development of a comprehensive set of tools for genome engineering in a cold- and thermo-tolerant *Kluyveromyces marxianus* yeast strain. Sci. Rep. 7, (2017).

38. Abdel-Banat, B. M. A., Nonklang, S., Hoshida, H. & Akada, R. Random and targeted gene integrations through the control of non-homologous end joining in the yeast *Kluyveromyces marxianus*. Yeast 27, 29–39 (2010).

39. Wilson, T. E., Grawunder, U. & Lieber, M. R. Yeast DNA ligase IV mediates non-homologous DNA end joining. Nat. 1997 3886641 **388**, 495–498 (1997).

40. Willmore, E. et al. A novel DNA-dependent protein kinase inhibitor, NU7026, potentiates the cytotoxicity of topoisomerase II poisons used in the treatment of leukemia. Blood 103, 4659–4665 (2004).

41. Yang, H., et al. Methods favoring homology-directed repair choice in response to CRISPR/Cas9 induced-double strand breaks. Int. J. Mol. Sci. 21, 6461 (2020).

42. Joberty, G. et al. A tandem guide RNA-based strategy for efficient CRISPR gene editing of cell populations with low heterogeneity of edited alleles. CRISPR. J. 3, 123 (2020).

43. Acosta, S., Fiore, L., Carota, I. A. & Oliver, G. Use of two gRNAs for CRISPR/Cas9 improves bi-allelic homologous recombination efficiency in mouse embryonic stem cells. Genesis 56, e23212 (2018).

44. Lane, M. M. et al. Physiological and metabolic diversity in the yeast *Kluyveromyces marxianus*. *Antonie van Leeuwenhoek*, Int. J. Gen. Mol. Microbiol. 100, 507–519 (2011).

45. Llorente, B. et al. Genomic exploration of the hemiascomycetous yeasts: 12. Kluyveromyces marxianus var. marxianus. FEBS Lett. 487, 71–75 (2000).

46. Barsoum, E., Martinez, P. & Åström, S. U. α3, a transposable element that promotes host sexual reproduction. Genes Dev. 24, 33 (2010).

47. Rajaei, N., Chiruvella, K. K., Lin, F. & Åström, S. U. Domesticated transposase Kat1 and its fossil imprints induce sexual differentiation in yeast. Proc. Natl. Acad. Sci. U. S. A. 111, 15491–15496 (2014).

48. Wu, L. et al. A protocol of rapid laboratory evolution by genome shuffling in *Kluyveromyces marxianus*. MethodsX 7, (2020).

49. Hickman, M. A. et al. The ‘obligate diploid’ *Candida albicans* forms mating-competent haploids. Nature 494, 55 (2013).

50. Fonseca, G. G., Gombert, A. K., Heinzle, E. & Wittmann, C. Physiology of the yeast *Kluyveromyces marxianus* during batch and chemostat cultures with glucose as the sole carbon source. FEMS Yeast Res. 7, 422–435 (2007).

51. Dubencovs, K. et al. Optimization of synthetic media composition for *Kluyveromyces marxianus* fed-batch cultivation. Fermentation 7, 1–21 (2021).

52. Walker, G. M. Metals in yeast fermentation processes. Adv. Appl. Microbiol. 54, 197–229 (2004).

53. Löser, C., Urit, T., Gruner, E. & Bley, T. Efficient growth of *Kluyveromyces marxianus* biomass used as a biocatalyst in the sustainable production of ethyl acetate. Energy. Sustain. Soc. 5, 2 (2015).

54. Hensing, M. C. M. et al. Effects of cultivation conditions on the production of heterologous α-galactosidase by *Kluyveromyces lactis*. Appl. Microbiol. Biotechnol. 43, 58–64 (1995).

55. Rech, R., Cassini, C. F., Secchi, A. & Ayub, M. A. Z. Utilization of protein-hydrolyzed cheese whey for production of β-galactosidase by *Kluyveromyces marxianus*. J. Ind. Microbiol. Biotechnol. 23, 91–96 (1999).

56. Rouwenhorst, R. J., Visser, L. E., Van Der Baan, A. A., Scheffers, W. A. & Van Dijken, J. P. Production, distribution, and kinetic properties of inulinase in continuous cultures of *Kluyveromyces marxianus* CBS 6556. Appl. Environ. Microbiol. 54, 1131–1137 (1988).

57. Lertwattanasakul, N. et al. Genetic basis of the highly efficient yeast *Kluyveromyces marxianus*: Complete genome sequence and transcriptome analyses. Biotechnol. Biofuels 8, (2015).

58. Mukherjee, P. K. et al. Socio-economic sustainability with circular economy — An alternative approach. Sci. Total Environ. 904, 166630 (2023).

59. Gómez-Márquez, C. et al. Diploid genome assembly of *Kluyveromyces marxianus* NRRL Y-50883 (SLP1). G3 Genes|Genomes|Genetics 12, (2022).

60. Lane, M. M. & Morrissey, J. P. *Kluyveromyces marxianus*: A yeast emerging from its sister’s shadow. Fungal Biol. Rev. 24, 17–26 (2010).

61. Mercier, G. et al. A haploid-specific transcriptional response to irradiation in *Saccharomyces cerevisiae*. Nucleic Acids Res. 33, 6635–6643 (2005).

62. Galitski, T., Saldanha, A. J., Styles, C. A., Lander, E. S. & Fink, G. R. Ploidy regulation of gene expression. Science (80-. ). 285, 251–254 (1999).

63. Keren, L. et al. Promoters maintain their relative activity levels under different growth conditions. Mol. Syst. Biol. 9, 701 (2013).

64. Petzoldt, T. Growthrates: Estimate growth rates from experimental data. CRAN (2022).

65. Pyne, M. E. et al. An engineered Aro1 protein degradation approach for increased cis,cis-muconic acid biosynthesis in *Saccharomyces cerevisiae*. Appl. Environ. Microbiol. 84, (2018).

66. Pacific Biosciences. SMRT Link: Explore and Analyze Your Data With Confidence. https://www.pacb.com/wp-content/uploads/Product-Brochure-SMRT-Link-Explore-and-Analyze-Your-Data-with-Confidence.pdf (2020).

67. Smit, A., Hubley, R. & Green, P. RepeatMasker Open-4.0. http://www.repeatmasker.org (2015).

68. Hoff, K. J. & Stanke, M. Predicting genes in single genomes with AUGUSTUS. Curr. Protoc. Bioinforma. 65, (2019).

69. Buchfink, B., Reuter, K. & Drost, H. G. Sensitive protein alignments at tree-of-life scale using DIAMOND. Nat. Methods 2021 184 18, 366–368 (2021).

70. Engler, C., Kandzia, R. & Marillonnet, S. A one pot, one step, precision cloning method with high throughput capability. PLoS One 3, (2008).

71. Ryan, O. W. et al. Selection of chromosomal DNA libraries using a multiplex CRISPR system. Elife 3, 1–15 (2014).

72. Stemmer, M., Thumberger, T., Del Sol Keyer, M., Wittbrodt, J. & Mateo, J. L. CCTop: An intuitive, flexible and reliable CRISPR/Cas9 target prediction tool. PLoS One 10, e0124633 (2015).

73. Labuhn, M. et al. Refined sgRNA efficacy prediction improves large- and small-scale CRISPR–Cas9 applications. Nucleic Acids Res. 46, 1375 (2017).

74. Zhang, Y. et al. A gRNA-tRNA array for CRISPR-Cas9 based rapid multiplexed genome editing in *Saccharomyces cerevisiae*. Nat. Commun. 10, 1–10 (2019).

75. Dykstra, C. B., Pyne, M. E. & Martin, V. J. J. CRAPS: Chromosomal-repair-assisted pathway shuffling in yeast. ACS Synth. Biol. 12, 2578–2587 (2023).

76. Todd, R. T., Braverman, A. L. & Selmecki, A. Flow cytometry analysis of fungal ploidy. Curr. Protoc. Microbiol. 50, e58 (2018).

