## Supplementary material for "Sequencing of a dairy isolate unlocks *Kluyveromyces marxianus* as a host for lactose valorization": Thornbury et al Supplementary Figures

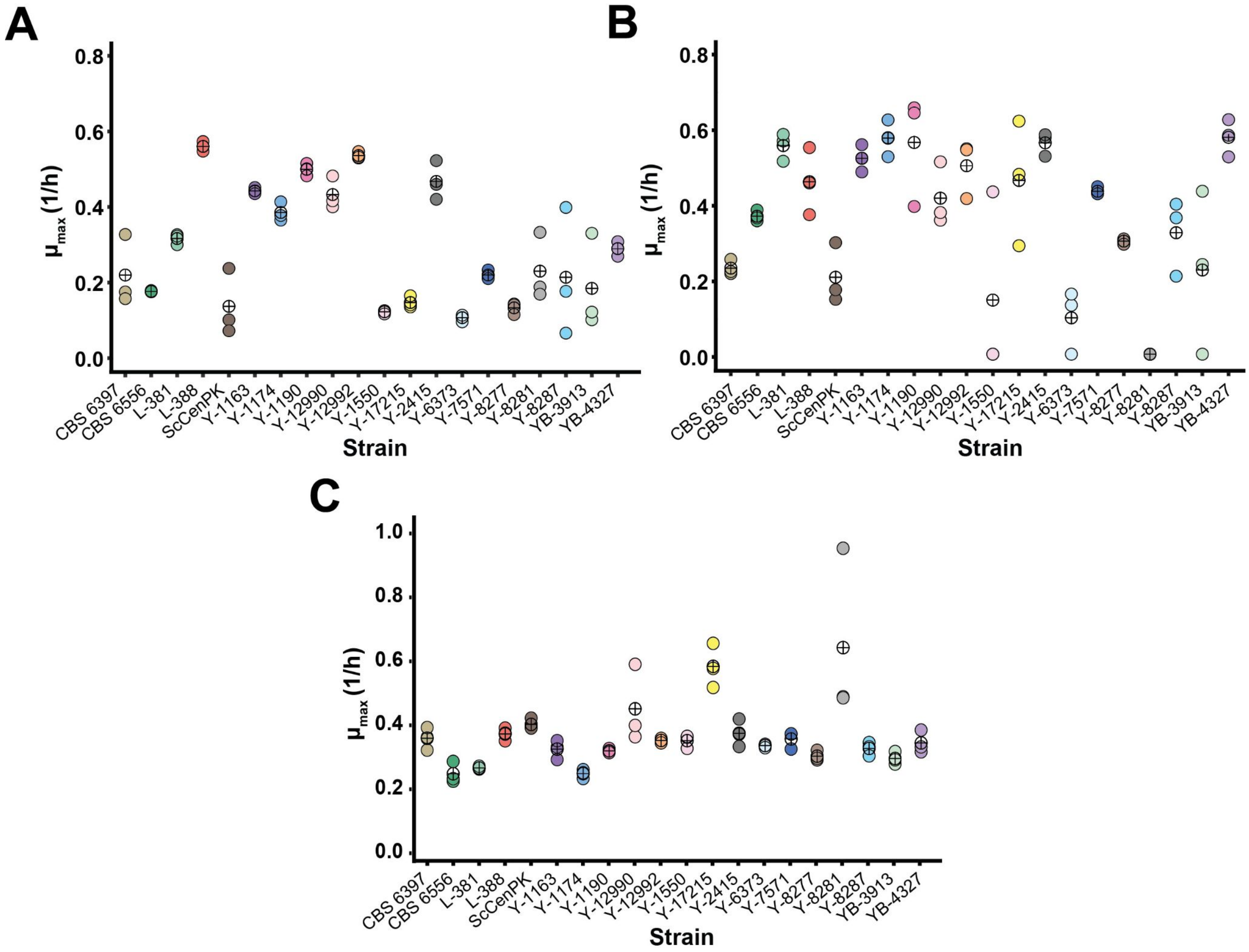


**Figure S1**. Growth rates of 19 strains of *K. marxianus* on **A**. YNBL, **B**. Supplemented permeate and **C**. YNBG. Each coloured dot represents a biological replicate, the + symbol represents the average of the three replicates.

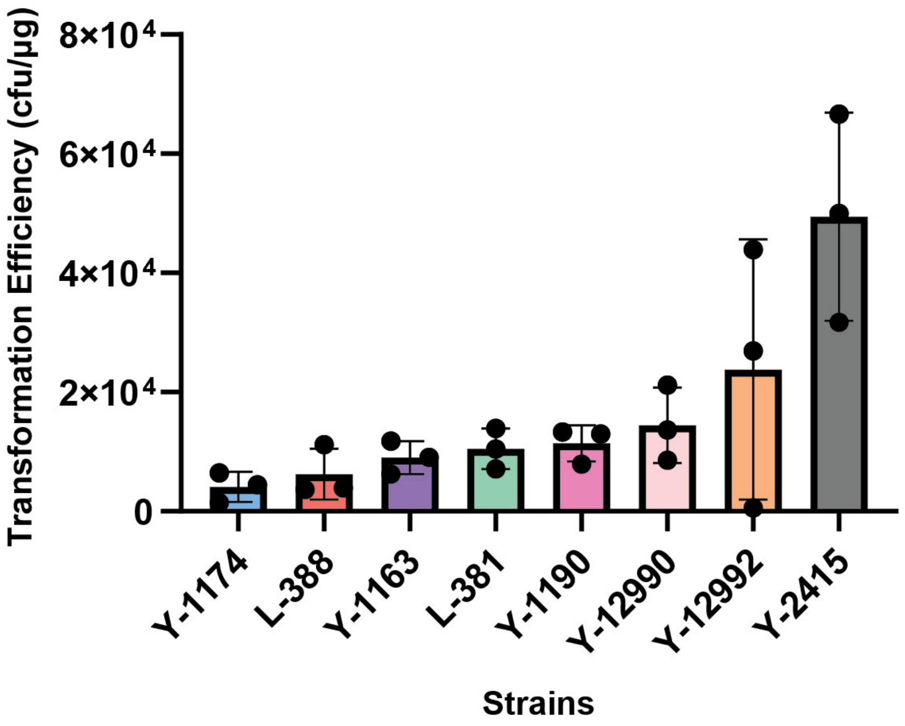


**Figure S2.** Transformation efficiency of our top growing strains of *K. marxianus* using a modified Gietz transformation method.

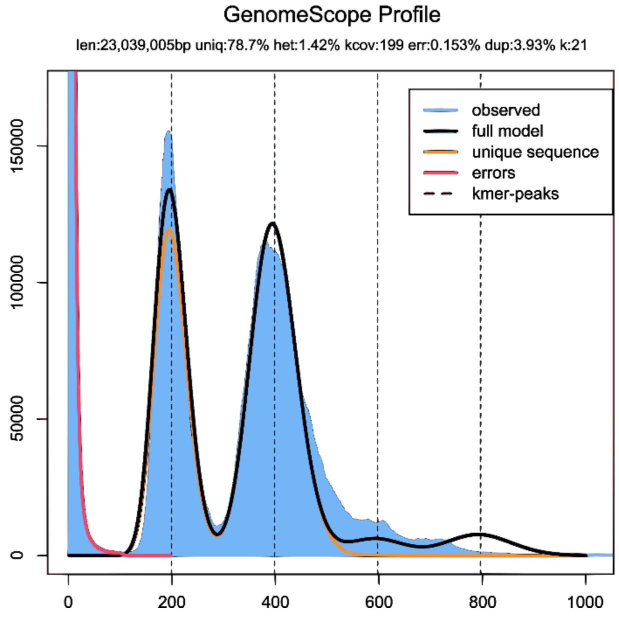


**Figure S3.** k-mer frequency of PacBio raw reads. k-mer coverage of Y-1190 using Jellyfish and Genome Scope.

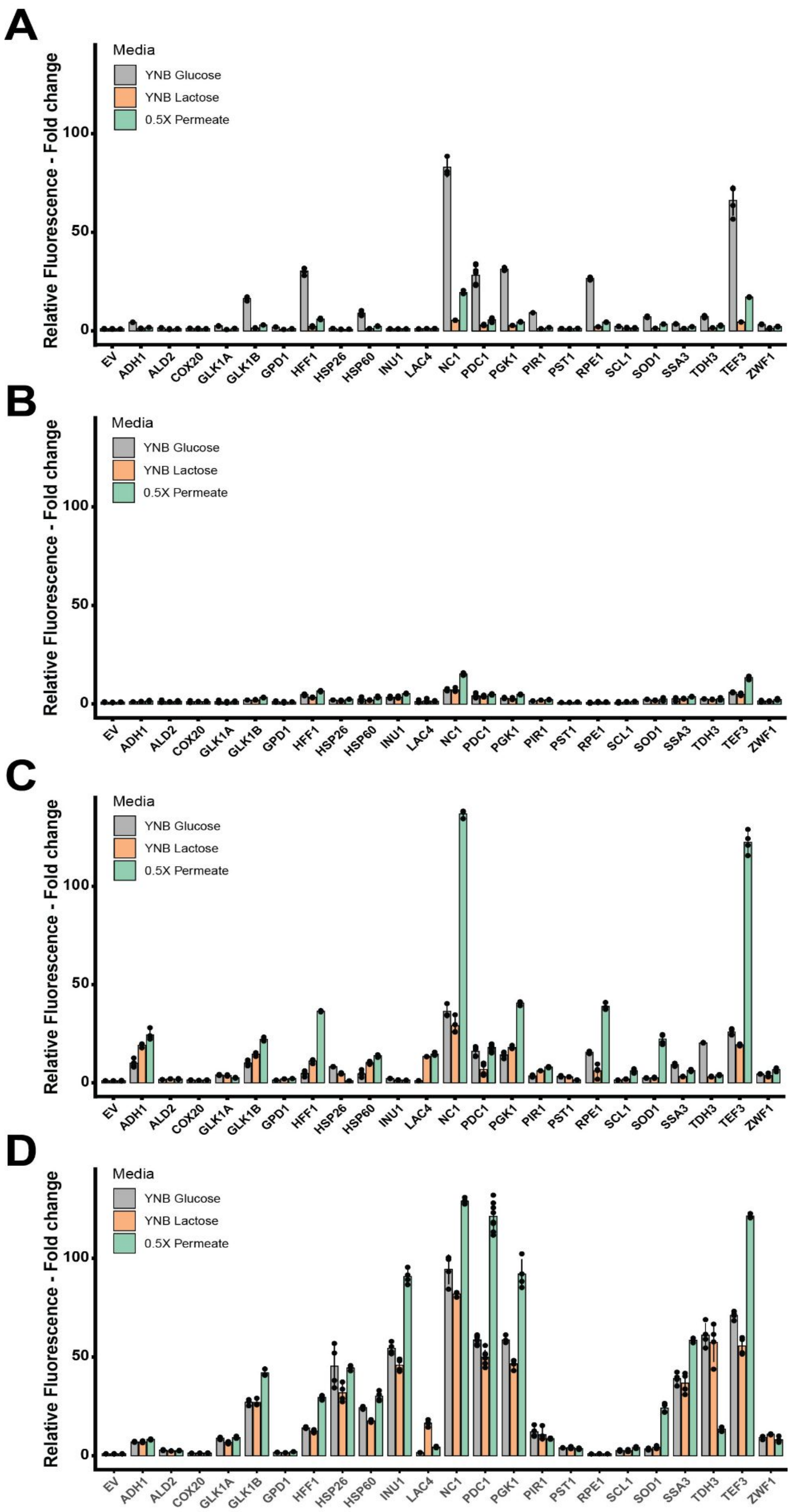


**Figure S4**. Relative promoter expression in *K. marxianus* **A**. CBS 6556 at 6 h, **B**. Y-1190 at 6 h, **C**. CBS 6556 at 24 h, **D**. Y-1190 at 24 h, YNBL, and supplemented permeate. Each promoter was tested in 4 biological replicates, the IQR method was used to remove outliers, and the error bars represent the standard deviation.

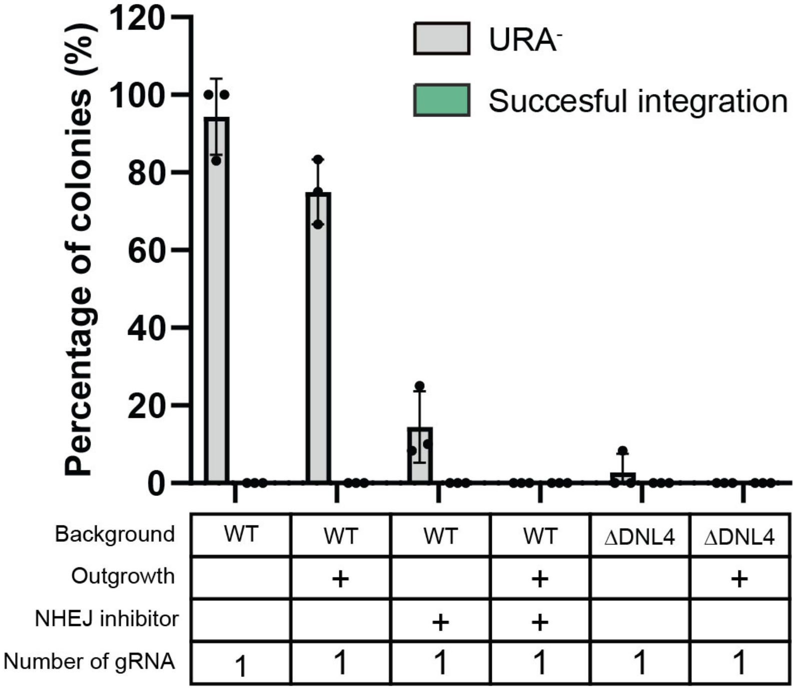


**Figure S5.** CRISPR-Cas9 genome editing efficiencies using one gRNA (sequence 2) for genomic deletion and integration via HDR. Each dot represents a biological replicate, and the error bars represent the standard deviation.

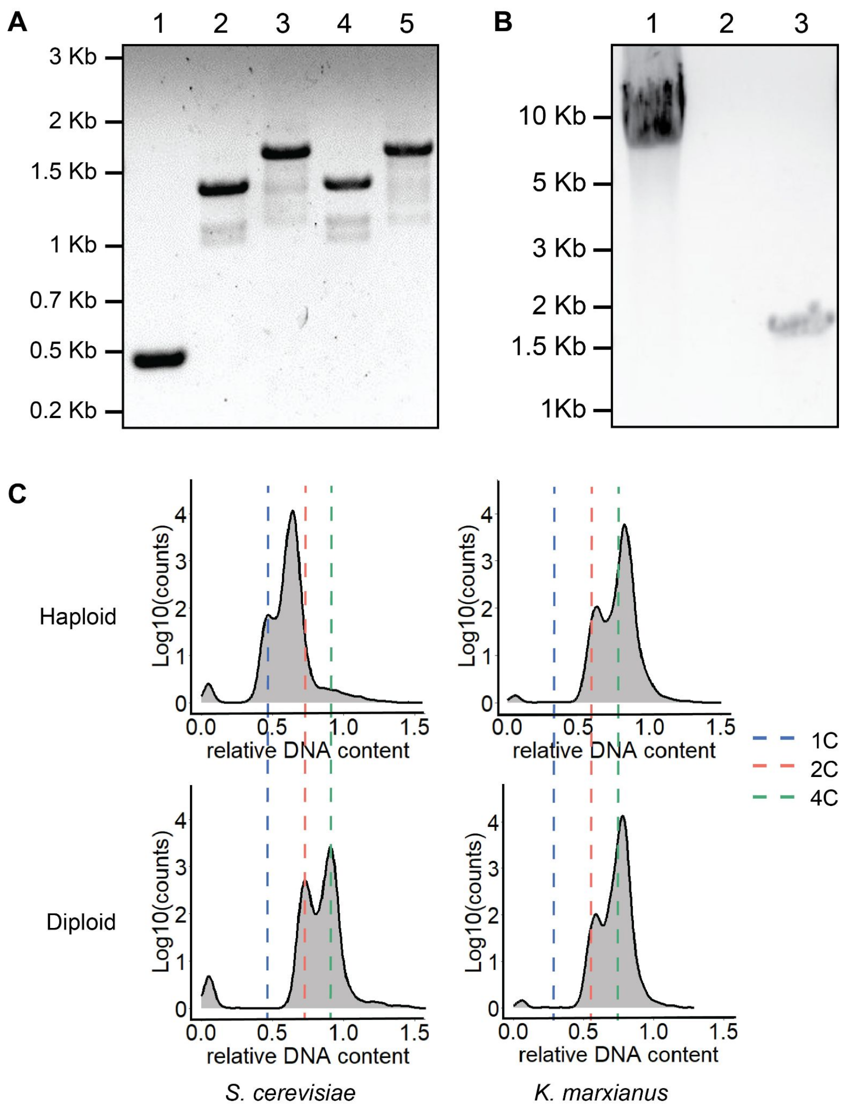


**Figure S6**. Ploidy and mating type analysis **A.** Detection of disrupted *Kat1* and *α3* in *K. marxianus* by PCR and agarose gel electrophoresis**.** Lane 1: *Kat1* amplification by primers MT169/MT172 (2kb in WT, 489bp when truncated). Lane 2: *α3* amplification from the *MAT* locus by primers JD108/JD110 (2.1kb in WT, 1.4kb when truncated). Lane 3: *α3* amplification from the *MAT* locus by primers JD088/JD109 (2.4kb in WT, 1.7kb when truncated). Lane 4: *α3* amplification from the *HML* locus by primers JD108/JD109 (2.1kb in WT, 1.4kb when truncated). Lane 5: *α3* amplification from the *HML* locus by primers JD088/JD109 (2.3kb in WT, 1.6kb when truncated). **B.** Determination of mating type following sporulation of *K. marxianus* Y-1190 (Δ*Kat1:*Δ*α3)*. Lane 1: Amplification of MAT locus using primers JD103/JD106 (6.8kb if MATα, 3.8kb if MATa). Lane 2: Amplification of MAT locus using primers MT240/MT241 (1.1kb if MATa). Lane 3: Amplification of MAT locus using primers MT240/MT242 (1.5kb if MATα). **C.** Determination of ploidy following sporulation of *K. marxianus* Y-1190 (Δ*Kat1:*Δ*α3)*. Propidium iodide staining followed by flow cytometry was performed to determine the ploidy of a known *S. cerevisiae* diploid (bottom left), haploid (top left), *K. marxianus* diploid (Y-1190 Δ*Kat1*:Δ*α3*, bottom right), and the sporulated MATα "haploid" (top right).

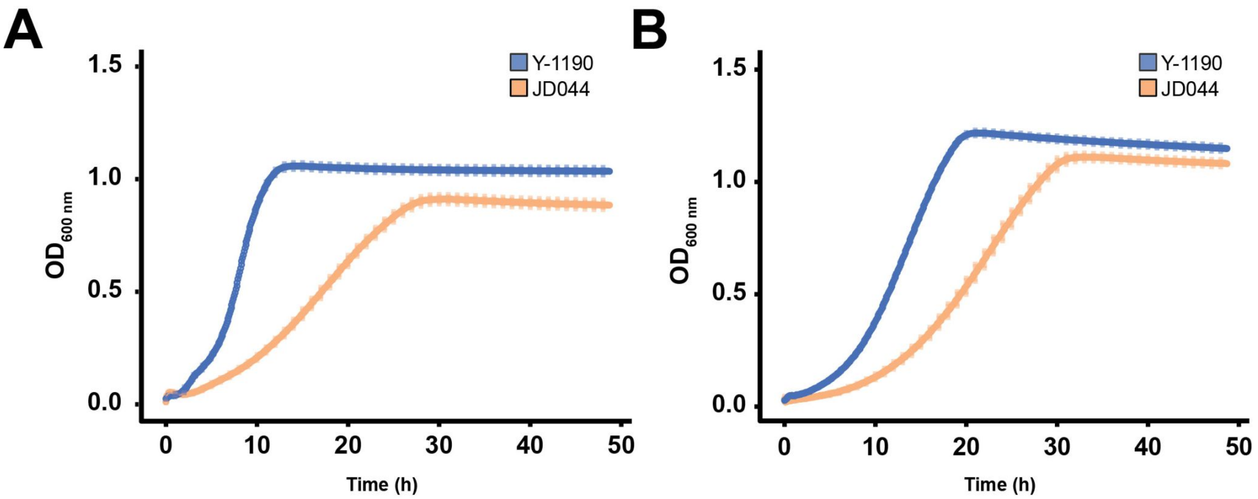


**Figure S7.** Growth curves of Y-1190 and homologous diploid JD044 in **A**. YNBL and **B**. Supplemented permeate. Growth was measured using $OD_{600}$ every 5 min for 48 hrs. The data points are an average of three biological replicates and the error bars represent standard deviation.

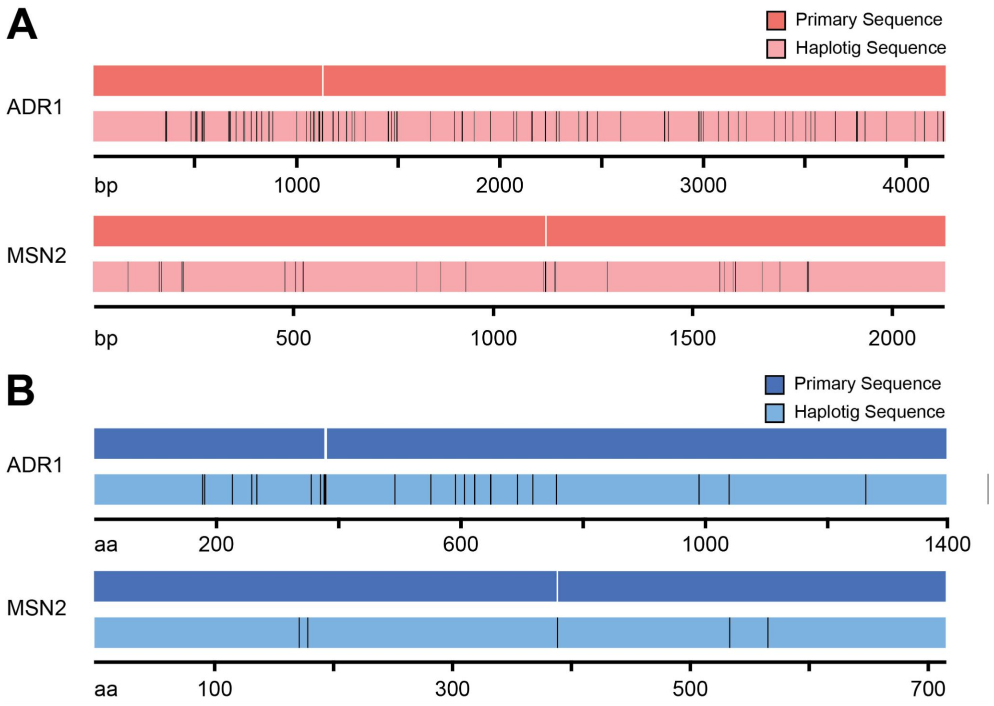


**Figure S8.** Primary and haplotig sequence comparison for two example genes **A.** Nucleotide sequences **B**. Amino acid sequences. Black lines represent changes in the A) nucleotide sequence and B) amino acid sequence.
